# Genomic epidemiology of vancomycin resistant *Enterococcus faecium* (VR*Efm*) in Latin America: Revisiting the global VRE population structure

**DOI:** 10.1101/842013

**Authors:** Rafael Rios, Jinnethe Reyes, Lina P. Carvajal, Sandra Rincon, Diana Panesso, Paul J. Planet, Aura M. Echeverri, An Dinh, Sergios-Orestis Kolokotronis, Apurva Narechania, Truc T. Tran, Jose M. Munita, Barbara E. Murray, Cesar A. Arias, Lorena Diaz

**Author notes:** Correspondence Lorena Diaz, PhD, Molecular Genetics and Antimicrobial Resistance Unit, International Center for Microbial Genomics, Universidad El Bosque, Av Cra 9 131a-02, Bogotá, Colombia.

## Abstract

The prevalence of vancomycin-resistant *Enterococcus faecium* varies across geographical regions yet little is known about its population structure in Latin America. Here, we provide a complete genomic characterization of 55 representative Latin American VR*Efm* recovered from 1998-2015 in 5 countries. We found that VR*Efm* population in the region is structured into two main clinical clades without geographical clustering. To place our regional findings in context, we reconstructed the global population structure of VR*Efm* by including 285 genomes from 36 countries from 1946-2017. Our results differ from previous studies showing an early branching of animal related isolates and a further split of clinical isolates into two sub-clades, all within clade A. The overall phylogenomic structure was highly dependent on recombination (54% of the genome) and the split between clades A and B is estimated to have occurred more than 3585 years BP. Furthermore, while the branching of animal isolates and clinical clades was predicted to have occur ∼894 years BP, our molecular clock calculations suggest that the split within the clinical clade occurred around ∼371 years BP. By including isolates from Latin America, we present novel insights into the population structure of VR*Efm* and revisit the evolution of this pathogen.

## Introduction

Enterococci are predominantly non-pathogenic gastrointestinal commensal bacteria that occasionally cause human infections. Among them, *Enterococcus faecalis* and *Enterococcus faecium* represent the species that account for most clinically relevant infections. In particular, *E. faecium* has been able to adapt to the hospital environment, emerging during the last few decades as a leading cause of health-care infections worldwide, and becoming the most challenging species to treat^1,2^.

Genome plasticity, the presence of multiple antibiotic resistance determinants, and the lack of therapeutic options have contributed to the adaptation of *E. faecium* to hospital environments^3,4^. Moreover, high recombination rates and the acquisition of mobile elements in the genome of *E. faecium* also have driven this evolutionary process^5^. In addition, the enrichment of virulence determinants, such as surface proteins and phosphotransferase systems (particularly PTS^clin^, a putative factor found to contribute to the intestinal colonization in a murine model) seems to provide an advantage to the hospital adaptive process^3,6^. Furthermore, functional gene groups, such as those involved in galactosamine metabolism, bile hydrolysis and phosphorus utilization, are also abundant in *E. faecium* clinical strains compared to non-clinical isolates, suggesting that specific metabolic factors have also facilitated adaptation^7^.

In terms of antibiotic resistance, one of the most relevant antibiotic resistance traits acquired by enterococci is resistance to vancomycin due to the *van* gene clusters^8^. Furthermore, vancomycin-resistant *E. faecium* (VR*Efm*) frequently exhibits resistance to ampicillin and high-level resistance to aminoglycosides^9,10^. Indeed, the World Health Organization (WHO) has categorized VR*Efm* as a priority agent for which the finding of new and effective therapeutic strategies is imperative^11^. VR*Efm* is widely distributed in hospitals around the world, with the prevalence varying according to geographical location. In US hospitals, VR*Efm* is an important clinical pathogen, particularly in immunosuppressed and critically-ill patients^1,12^. The National Health-Care Safety Network described that 82% of *E. faecium* recovered from bloodstream infections in the US were vancomycin-resistant, whereas only 9.8% of *E. faecalis* were resistant to vancomycin^12^. In Europe, prevalence rates of VR*Efm* vary widely by country, but according to the European Centre for Disease and Control (ECDC) 2016 report, overall prevalence across European countries was 30%^13^. Although data regarding VR*Efm* in Latin America are scarce, a few studies have shed some light on the current situation. A prospective multicentre study focusing on 4 countries in northern South America (i.e. Colombia, Ecuador, Peru and Venezuela) found an overall prevalence of VR*Efm* in clinical enterococcal isolates of 31%^14^. More recently, another study performed in Brazil reported a VR*Efm* prevalence close to 60%^15^.

Tracking the population structure of *E. faecium* using conventional bacterial typing techniques has been challenging^16^. Although wide genetic variability has been observed among *E. faecium* strains causing clinical infections, a previously described lineage (designated clonal complex CC17 by multi locus sequence typing [MLST]), was initially recognized as globally distributed^17^. However, the classification of this lineage by MLST has some important drawbacks when analysing the population structure of *E. faecium*, since high rates of recombination in the MLST loci often occurs in these organisms^18^. Additionally, some strains are not type able by MLST due to the lack of the locus *pts*^19^ leading to major discrepancies compared to whole-genome sequencing (WGS) when it is used for typing purposes^20^.

Whole-genome-based comparative phylogenomic analyses using *E. faecium* recovered from different geographical regions have identified two clades, designated A and B. Clade A mostly contains isolates recovered in clinical settings (including those from CC17)^21^, while clade B encompasses organisms isolated in community settings, usually from healthy individuals ^3,20,22–24^. A further subdivision has been described within clade A, which groups isolates from animal origin in a subclade (designated as A2), separating them from those recovered from human infections or colonization (subclade A1).

However, these analyses have been performed mostly with US and European isolates, lacking geographical diversity particularly in areas such as Latin America. Indeed, studies on the molecular epidemiology of VR*Efm* isolates from Latin America are sparse, with one study suggesting that the CC17 lineage predominates^14^. Furthermore, studies analysing the population structure of VR*Efm* in the region using high-resolution, WGS-based phylogenomic comparative methods are limited. Here, we sought to characterize the population structure of VR*Efm* lineages in a collection of isolates recovered between 1998-2015 in prospective multicentre studies performed in selected Latin-American hospitals^14,25,26^ and revisit the global population structure and evolutionary history of VR*Efm*.

## Results

### Genomic characterization of Latin American VR*Efm* clinical isolates

From a collection of 207 VR*Efm* clinical isolates obtained between 1998 and 2015 in five Latin American countries (Colombia, Ecuador, Venezuela, Peru and Mexico), we selected 55 representative isolates for WGS. We included the first VR*Efm* (ERV1) reported in Colombia as the representative of 23 isolates with identical PFGE banding pattern, recovered from an outbreak in 1998-1999 and affecting 23 patients in a single teaching hospital^25^. Five isolates (out of 7 available) were selected from a national surveillance in Colombia during 2001-2002, which included 15 tertiary hospitals among 5 cities^26^ and 16 (out of 35 available) were chosen from a subsequent surveillance study (2006-2008) performed in Colombia, Ecuador, Venezuela and Peru and the selected isolates were chosen based on their different banding patterns^14^. The remaining 33 isolates were obtained from sporadic isolates and outbreaks that occurred in Colombia and Mexico (2002-2014). In order to characterize the VR*Efm* lineages circulating in Latin America, we reconstructed their phylogenetic history based on 1,674 genes (groups of orthologous sequences; hereafter referred to as orthogroups) present in more than 90% of the genome sequences (core genome) from a total of 6735 orthogroups (pan-genome) using a Bayesian approach (Figure 1A). We observed a split into two main clades (Clade I and Clade II, marked in red and green, respectively). Clade I included all the ST412 isolates, while Clade II had all the ST17 isolates from our sample. We observe that the emergence of VR*Efm* in Colombia was associated with Clade II, including the first VR*Efm* (described in 1998) and representatives from the first national surveillance (2001 to 2002). Additionally, ST412 was reported in 2005 and, since then, ST17 and ST412 seem to be the most prevalent STs in the country. Our previous results showed that Peru had the highest prevalence of VRE*fm* (48%) and our PFGE and MLST results suggested higher diversity in Peruvian lineages compared to Colombia, Ecuador and Venezuela with a predominant circulation of ST412^14^. Indeed, the representative VR*Efm* isolates of the circulating lineages in Peru collected in the two-year period (2006-2007) exhibited a marked genomic variability (Figure 1A and B).

**Figure 1.**
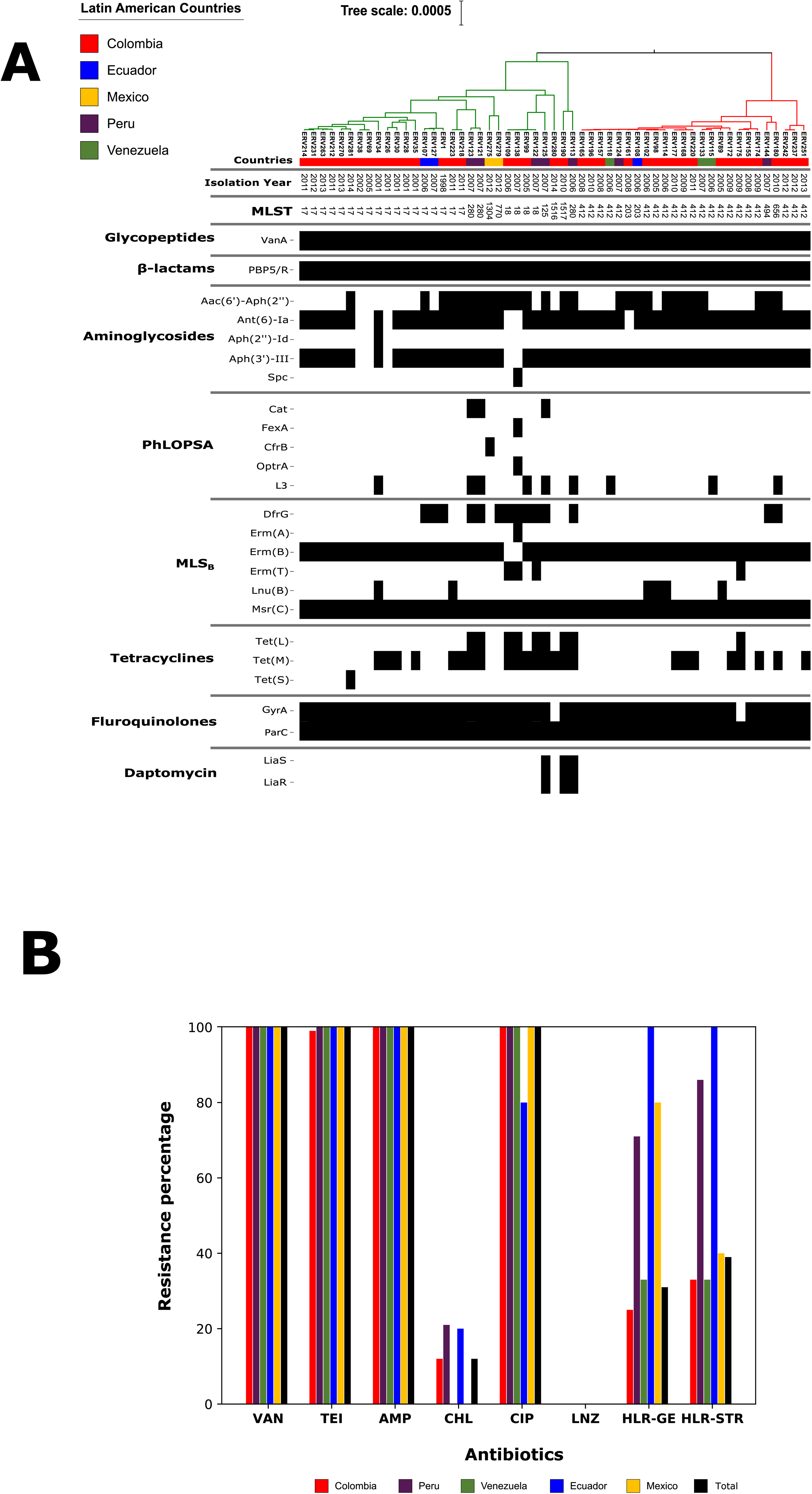
(A) Bayesian phylogenomic tree from the core genome and genomic characterization of resistance elements of 55 representative Latin American VR*Efm* strains; the presence of a genetic element is marked as a black box in the corresponding column of the isolate. (B) Phenotypic resistance profile of 207 clinical isolates of VR*Efm* from our Latin American collection for vancomycin (VAN), teicoplanin (TEI), ampicillin (AMP), chloramphenicol (CHL), ciprofloxacin (CIP), linezolid (LNZ), high-level resistance to gentamicin (HLR-GE) and high-level resistance to streptomycin (HLR-STR).

### The resistome and virulome of Latin American VR*Efm*

In order to characterize antibiotic resistance determinants, we built resistome profiles by detecting acquired resistance genes and mutations known to confer resistance to linezolid, ciprofloxacin and daptomycin. All the VR*Efm* isolates from our collection were resistant to vancomycin (MIC_90_ >256 µg/ml) and teicoplanin (MIC_90_ 64 µg/ml) (Figure 1B). The presence of *vanA* was confirmed in all isolates by PCR assays. Consistently, we confirmed the presence of the entire *vanA* cluster in 54 out of the 55 sequenced genomes. Of note, the genome of ERV69 lacked the two-component regulatory system *vanSR*, although still exhibiting MICs of >256 µg/ml and 64 µg/ml for vancomycin and teicoplanin, respectively. The deletion of the two-component regulatory system has been previously reported^27^.

High-level resistance to ampicillin was consistently found in all 55 *E. faecium* isolates, a phenotype that was corroborated using comparisons of the PBP5 protein sequence using a machine-learning prediction model. This approach was based on the amino acid changes present in the PBP5 protein across susceptible and resistant isolates (see details in Methods).

High-level resistance to gentamicin was identified in 31% of the isolates of our collection and, within the sequenced representatives, the presence of *aac(6’)-aph(2”)* was detected in 49% of the genome sequences. High-level resistance to streptomycin was identified in 39% of the Latin American VRE*fm* isolates with a high prevalence of the *ant(6)-Ia* gene (89%; n=49) in the sequenced genomes.

Fluoroquinolone resistance is very common in *E. faecium*. All isolates in our collection were fluoroquinolone-resistant and we were able to predict the presence of substitutions in GyrA and ParC associated with this phenotype. The most common substitution in GyrA was Ser84Arg (67%; n=37). All isolates exhibited Ser82Arg (53%; n=29) or Ser82Ile (47%; n=26) substitutions in ParC.

The *cat* gene conferring resistance to chloramphenicol was present only in the three Peruvian genomes. Interestingly, Peruvian isolates had the highest resistance to this antibiotic (21%). All the isolates from this collection were susceptible to linezolid; however, we detected the various genetic elements previously associated with linezolid resistance. The gene, *optrA*, was detected in one genome of a Colombian linezolid-susceptible isolate (ERV138). Also, we identified the presence of *cfrB*, a recently reported variant of *cfr*^28^, in a Mexican isolate (ERV275). We predicted tetracycline resistance owing to the presence of *tetM* (43.6%; n=24), *tetL* (16.3%; n=9) and *tetS* (1.8%; n=1) in the sequenced genomes, but resistance to this group of antibiotics was not tested phenotypically. Substitutions in LiaS (Thr120Ala) and LiaR (Trp73Cys), which have been strongly associated with daptomycin resistance and tolerance^29,30^, were present in three VR*Efm* isolates, recovered before daptomycin was available in the region. Of note, the three isolates showed MICs between 2-4 µg/ml, considered now as “daptomycin-susceptible dose-dependent”, by the Clinical & Laboratory Standards Institute (CLSI)^31^.

Latin American VRE isolates also harboured a high proportion of putative virulence determinants (Figure 2). The vast majority had gene clusters related to pilus formation, adhesins and microbial surface components recognizing adhesive matrix molecules (MSCRAMMS). Interestingly, the notable exception was the Clade I isolates, which often lacked *fms22, swpC* and *hyl_Efm_*. Our results suggest that the “virulome” of Latin-American VRE is similar to those from other regions in the world^32^.

**Figure 2.**
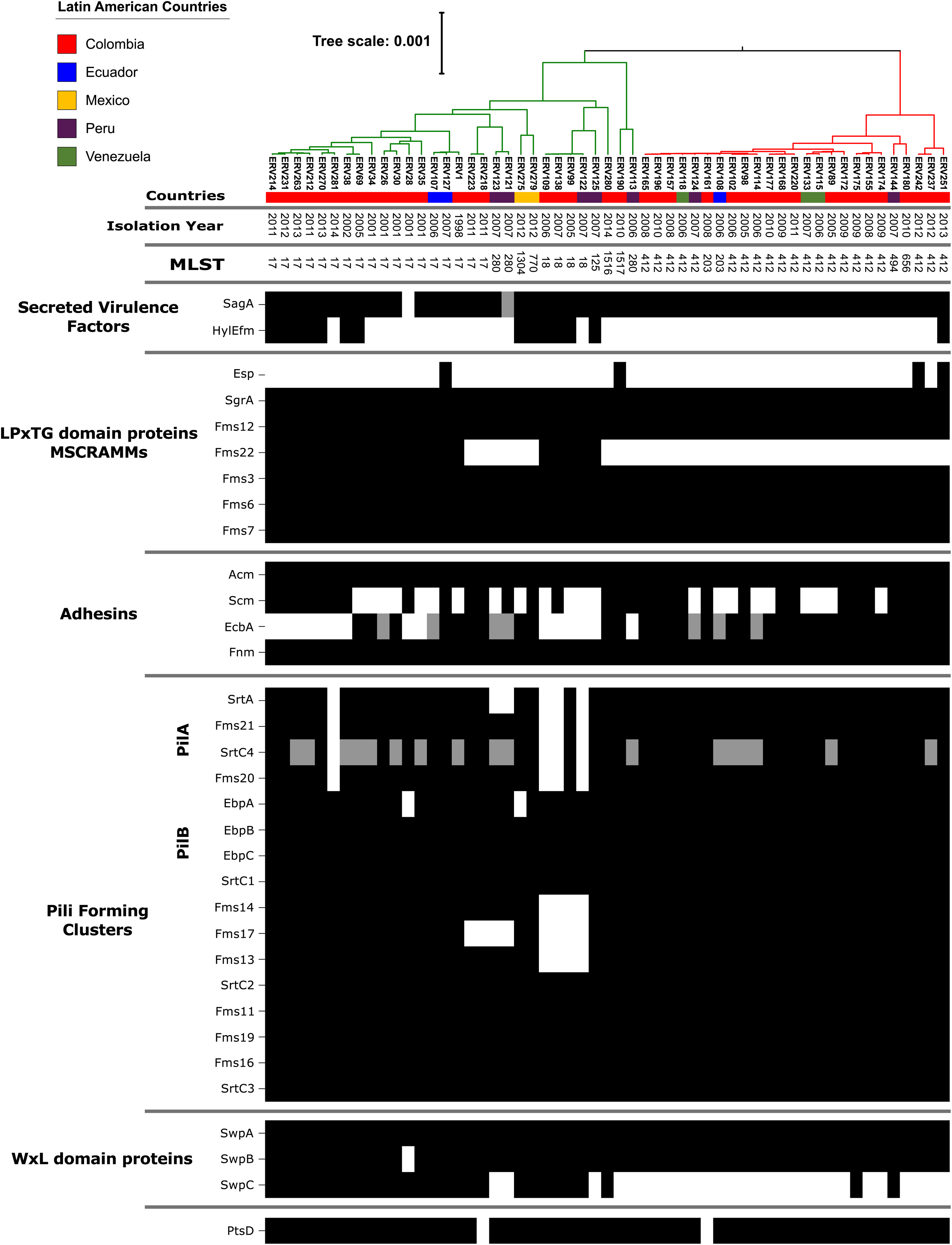
Bayesian phylogenomic tree from the core genome and genomic characterization of virulence factors of 55 representative Latin American VR*Efm* strains, the presence of a genetic element is marked as a black box in the corresponding column of the isolate.

### Global Phylogenetic Reconstructions of Latin American VRE

To place the genetic lineages of VR*Efm* isolates circulating in Latin America into a global context, we performed a WGS-based phylogenomic analysis. We included 285 *E. faecium* genomes (VRE and non-VRE) from the publicly available NCBI collection aiming to incorporate a diverse set of sequences for comparisons. The included isolates were from colonizing, commensal, animal and clinical sources and were collected between 1946-2017 from Europe, North America, Asia, Africa and Australia (Supplementary Table 1). We constructed a pangenome (29,503 orthogroups) and core genome (978 orthogroups). Using the core genome, we built a phylogenomic tree of the species to show the evolutionary relationships among isolates based on the variation of their genomic sequences. Figure 3 shows that, as previously reported, we found a clear split into two main clades corresponding to the previously designated clades A and B^3,22,24^. All Latin American isolates from our clinical collection were in clade A. We compared the genomic characteristics among the two main clades and found similar findings as published previously (Supplementary table 2)^3^. However, our data showed that the core genome was larger in clade B as compared to clade A (1,466 vs 1,182 orthogroups, respectively).

**Figure 3.**
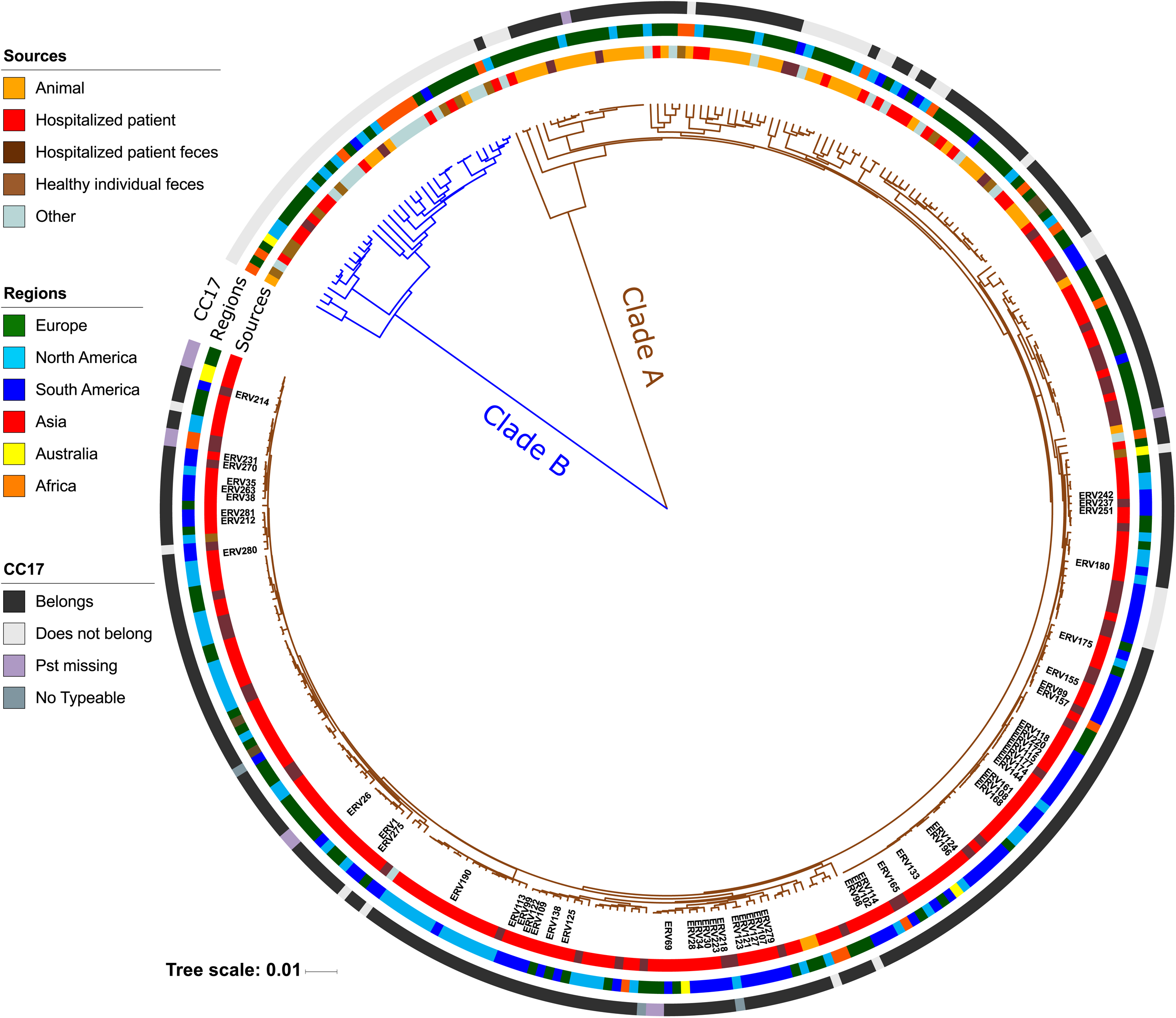
Bayesian phylogenomic tree from the core genome of 340 genomes sampled from 36 countries between 1946 and 2017 and from different sources. Blue branches showed the genomes grouped within clade B, while brown branches show isolates from clade A. The outer coloured rings (from inner to outer) indicate the source of each isolate, the region from which it was sampled and its relationship through MLST typing (if possible) to Clonal Complex 17. Labels show the isolates originating from our Latin American collection.

Considering the relevance of *E. faecium* as a cause of hospital-associated infections and that all Latin American isolates were grouped within clade A, we sought to dissect the population structure of this clade when adding the genomes of these isolates. Our first approach was based on a core genome (>90% reconstruction), which contained 1,226 orthogroups and the isolate Com15, from clade B, as the outgroup to root the tree. We observed two major subclades. The first was composed of 52 genomes, most of which were from animal sources (57%, n=30), related to the previously described subclade A2^3^. The second lineage harboured 273 genomes, with 91% (n=228) corresponding to isolates obtained from clinical sources (Supp. Figure 2A), and related to the subclade A1^3^.

Previous studies have shown contradictory distributions of the subclades A1 and A2 within clade A^20^; suggesting that clade A2 is not in fact a clade, but rather corresponds to the paraphyletic early branching lineages of clade A. To further clarify the issue, we performed a phylogenomic analysis accounting for recombination events within clade A. We used the variants found from paired alignments of each genome against the chromosome of reference Aus0085 and built a whole-genome multiple sequence alignment (WGMSA) of all genomes in the clade. We used this alignment to create a maximum likelihood tree, which is required for determining recombinant regions using ClonalFrameML^33^. The average amount of recombination found in the 303 genomes belonging to clade A was 19,539pb (Supp. figure 2C). The total recombinant regions found across clinical isolates encompassed 1.6 Mb (54% of the length of WGMSA). Interestingly, the exclusion of recombinant regions considerably altered the structure of the tree, and showed 7 early-branching subclades that included 73 genomes (mostly from animal sources) rather than a split into clades A1 and A2.

Following these animal-related early branches, we observed a split into two main subclades (Supp. Figure 2B). Overall, these subclades were related to clinical sources, exhibiting a high similarity in terms of prevalence of antibiotic resistance and virulence determinants (Supplementary table 3). We refer to them as clinically-related clades I and II (CRS-I and CRS-II), containing 101 and 124 genomes respectively. Latin American genomes from our collection were split between these two CRS, showing that Clade I and Clade II (derived from the analysis of Latin American VR*Efm*, see above) belonged to CRS-I and CRS-II, respectively. Of note, the genomes from our collection were distributed almost equally between CRS-I (49%) and CRS-II (51%). Furthermore, despite the inclusion of a few outbreak isolates and that VR*Efm* from Latin America originated in different periods, cities and countries, our phylogenetic reconstruction showed 11 conserved clusters with four or more isolates from the same country (Figure 4). In particular, three clusters had only Colombian genomes with the number of SNPs among them, within the regions not showing recombination, ranging between 36 and 160. We also found clusters among isolates from Brazil (n=3), USA (n=3), Denmark (n=1) and Sweden (n=1). The Danish cluster is situated in the animal-associated branches, and their genomes were closely related (with an average of 43 SNPs among them). Of note, two of the USA clusters were related to each other and to 5 other isolates, four of them from the UK and one from Colombia in our collection (172 SNPs difference on average among them).

**Figure 4.**
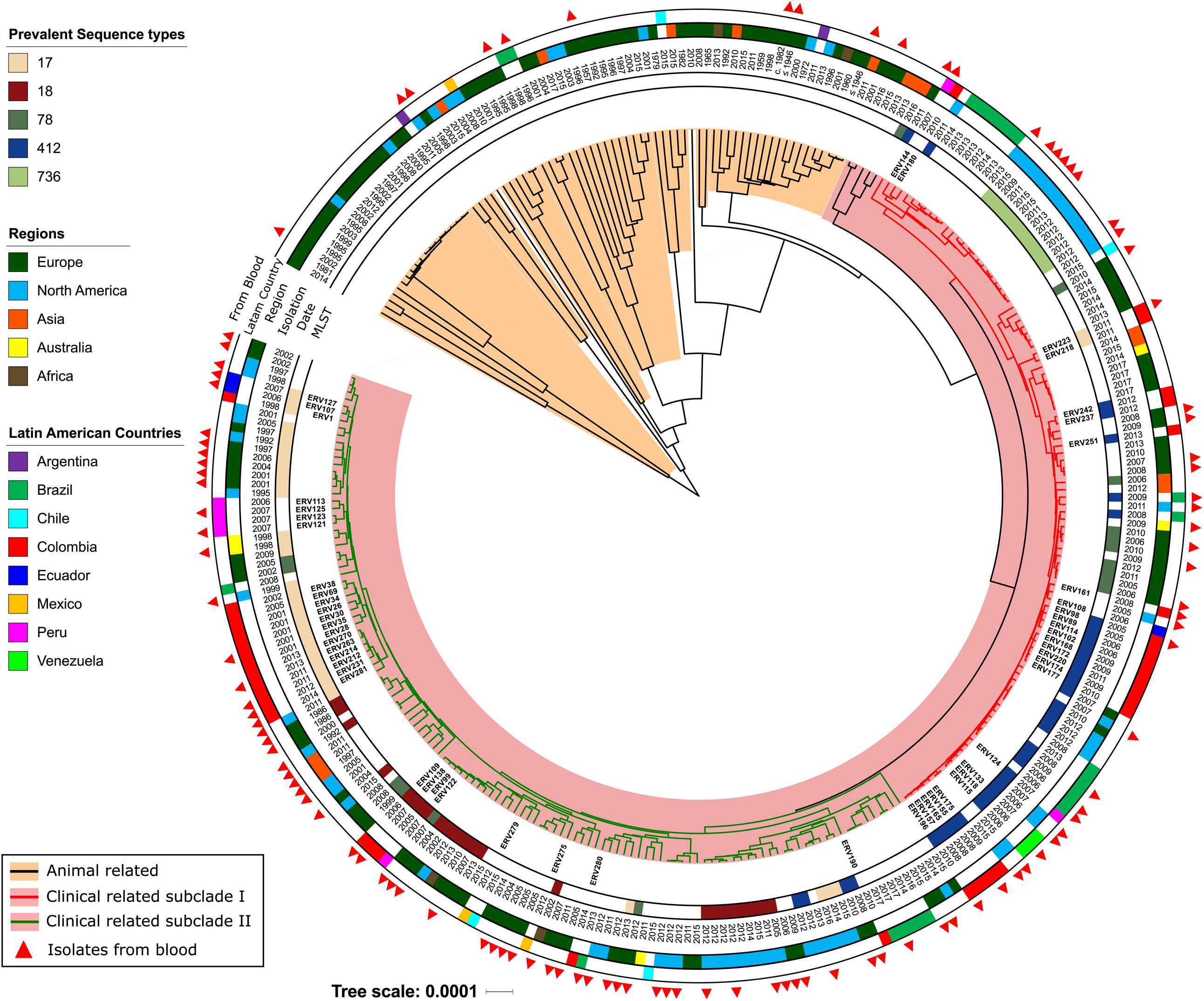
Bayesian phylogenomic tree from the non-recombinant regions of the 303 Clade A genomes. Branches highlighted in orange represent genomes from the animal early branches. Branches highlighted in pink show genomes from clinical related isolates. Red and green branches show the genomes from clinically related subclades (CRS) I and II, respectively. Annotation rings (from inner to outer) show the sequence type (ST) of the isolate (only the five most prevalent STs in the sample are shown), the isolation year, the region from which the isolate was sampled and if the region was Latin America, the exact country from where it was obtained. The last ring shows which isolates were recovered from blood.

In CRS-I, there were 23 different STs, with ST412 and 78 the most frequent (34% and 11%, respectively) (Figure 4). Importantly, we did not find a strong correlation between MLST and the phylogenomic analysis, as isolates belonging to the same ST were not all clustered in the same clades, and were distributed in different groups in the phylogeny. In particular, 56% (n=9) of genomes from ST78 were in CRS-I, while 37% (n=6) were in CRS-II. To further dissect this discrepancy, we performed a phylogenetic reconstruction using only the sequences of the 7 MLST *loci* and compared it against the phylogeny of Clade A. Our results showed that many isolates from ST17, ST18, ST78, ST203, ST412 were in different clusters and even formed subclades in the non-recombination reconstruction (Supplementary Figure 3).

In relation to antibiotic resistance determinants, we found important differences comparing the presence/absence of genomic elements associated with antibiotic resistance between the CRSs and the animal branches. Indeed, the animal-associated branches exhibited a lower prevalence of elements associated with glycopeptide (34.2%), aminoglycoside (21.9%), ampicillin (9.5%) and fluoroquinolone resistance (2.7%) compared to the CRS isolates, which harboured these determinants in 78%, 85%, 95% and 99% of isolates, respectively. In contrast, similar frequencies of determinants coding for resistance to macrolides (>98%), tetracyclines (between 50-63%) and oxazolidinones (between 2-12%) were found between animal and clinical clades (Supp. Table 3). Within the subclades of clade A, only 9% of isolates within the animal-associated branches exhibited resistance to ampicillin (7 out of 71 complete PBP5 sequences), while 99% of the clinically related subclades (100% in CRS-I and 98% in CRS-II) harboured the predicted *pbp5-R* allele^34,35^. Mutations associated with fluoroquinolone resistance were also much more highly prevalent in clinical clades (>98% for CRSs) vs animal branches (2.7%; p<0.001).

Genes encoding putative surface adhesin proteins (e.g., *acm, scm, esp, sgrA, fms6* and *fms22*) and two of the pilus-forming clusters were significantly more common in the CRSs, (p-values below 0.001 in all cases) compared to animal isolates (Supp. Table 3). We next compared the presence/absence of putative mobile elements between animal branches vs. CRSs. On average, the number of insertion sequences in the former were 5.7, whereas the clinical subclades had 6.9 (6.76 CRS-I and 7.06 for CRS-II). Of note, *rep17* was notoriously overrepresented in the CRSs (Supp. Table 3), located in the plasmid pRUM, which is a representative member of rep17 family and has been associated with the toxin/antitoxin system Txe/Axe^36^.

### Rates of evolution across the whole population of *E. faecium*

Using the sampling date of isolates within clade A, we performed molecular clock analyses on the entire clade A and its subgroups (animal branches, CRS-I and CRS-II). We found that the oldest split within clade A likely occurred ∼3,585 years ago (y.a.) (95% High Posterior Density Interval [HPDI]: [2626, 4690]). The separation of the clinical subclades from the animal branches is predicted to have occurred ∼894 y.a. (95% HPDI: [649, 1171]) (Supplementary Figure 3). The most recent split between CRS-I and CRS-II was dated ∼371 y. a (95% HPDI: [272, 488]) (Supplementary Figure 4). The substitution rate across the clade A genomes was 3.91E-7 (95% HPDI: [2.78E-7, 5E-7]), which translates to 0.53 SNPs per year (using only non-recombinant regions or 1.17 SNPs if the WGSA is used). The substitution rates within each subgroup of genomes were 3.02E-7 (95% HPDI: [2,78E-7, 3,46E-7]) for animal branches, 4.7E-7 (95% HPDI: [4,01E-7, 4,98E-7]) for CRS-I and 4.63E-7 (95% HPDI: [3.92E-7, 4.98E-7]) for CRS-II. These rates are equivalent to 0.41, 0.64 and 0.63 SNPs per year for animal branches, CRS-I and CRS-II, respectively. Our results support that clinically related clades are evolving faster than those of the animal branches.

## Discussion

Our results indicate that VR*Efm* is widely present in Latin America but that their frequency and population structure seem to vary from country to country. As multicentre analyses of VR*Efm* in the Latin American region are rare, our study is unique in its dissection of the population structure of VR*Efm* in the region. Unlike previous studies, we found two distinct populations of clinically-related isolates of VR*Efm*. This subpopulation separation was also seen in our analyses of the global population of *E. faecium*. The causes for the splitting of the population structure of VRE (CRS-I and CRS-II) are not clear, but the findings were consistent when analysing the population structure in the presence or absence of recombinant regions. Such a separation suggests that these lineages have been expanding through Latin American countries and highlights the importance of establishing genomic surveillance studies for these multidrug-resistant organisms. Furthermore, the distribution of the Latin American isolates across the tree does not suggest a particular dominance of a specific lineage circulating in the region or country, suggesting that the presence of VR*Efm* in Latin America is associated with multiple introductions of VR*Efm* lineages that are circulating globally. Interestingly, some South American countries such as Brazil (no isolates available for this study) have reported VR*Efm* since 1997^37^, and their prevalence appears to be increasing exhibiting a shift from *E. faecalis* to VR*Efm* since 2007^15^. Of interest, ST412 isolates reported in some regions of Brazil^38,39^ have also been detected in some Caribbean countries^40^ and this sequence type was also identified in our collection in Colombia, Peru and Venezuela since 2005^14^, suggesting wide dissemination of this genetic lineage.

Our VR*Efm* phylogenomic analysis, which includes a highly diverse sample collection and excludes recombinant regions from the genome, questions the presence of a single animal clade. Our results suggest that the animal isolates represent multiple lineages that diverged prior to the emergence of the clinical subclades in the clade A^3^. Importantly, animal-associated branches have significantly lower predicted ampicillin resistance, fluoroquinolone resistance mutations, virulence elements and average number of insertion sequences, similarly to what has previously reported^41^. Furthermore, the amount of recombination in clade A genomes was greater than previous results. Importantly, this difference (54% vs 44% found in previous studies^18,42^) could be due to the fact that previous analyses were based on the alignment of SNPs from a core genome and neither included non-coding regions nor invariant sites to identify the recombinant DNA. Over the recombinant regions, we found partial sequences in 5 out of the 7 loci used by MLST (*ddl, gyd, purK, gdh* and *adk*), corroborating the notion that the current *E. faecium* MLST scheme has major limitations to describe the population structure of VR*Efm*. Interestingly, the exclusion of recombinant regions considerably altered the structure of the tree, dissolving the animal-related clade into a paraphyletic group and reducing the length of the branches across the tree (Supplementary Figure 2). Additionally, we found a lack of concordance between MLST classification and the clades. The discrepancy is likely explained by the presence of recombinant regions in the MLST genes, low variation in some of the loci, and the absence of *pst* in many isolates^19,20,43^

Previous studies estimated that the separation between clades A and B occurred 2776 ± 818 y.a.^3^, a time frame that is similar to our results. However, the previously reported split between animal branches and the clinically-related subclades was reported to occur 74 ± 30 y. a., which is much more recent than what we found. Our findings showed at least a tenfold lower mutation rate from what has been previously reported^3,18^. This finding could be associated with the larger genomic region used in our analysis and the increase in the diversity of the sampled genomes. Indeed, dating of the splits between the animal-associated branches and the clinically-related subclades, and the lower mutation rates across clade A correlates with lower number of SNPs per year. It has been estimated that the *Enterococcus* as a genus arose around 500 million years ago^44^ and ancient isolates of *E. faecium* have been found in permafrost over 20,000 y.a.^45^, supporting our findings that a more ancient branching between Clade A and B could have occurred.

Our study could be subject to sampling bias due to small sample size of genomes from Latin America, but we attempted to include as many and as diverse genomes as possible from our collection, based on phenotypic characteristics and PFGE typing of the strains. Also, we included all publicly available genomes from the region, provided that the associated demographic information was complete (source, year of sampling and geographical location), which also reiterates the low number of previously sequenced genomes of *E. faecium* in Latin America at the moment of sample selection. Nonetheless, our results supporting the existence of two clinical subclades were maintained even after the inclusion of genomes from other continents; that is, our conclusion holds beyond sample size, further indicating that the population structure of the clinical related isolates is divided into two main lineages within clade A.

## Conclusions

We provide comprehensive insights into the genomic epidemiology of VR*Efm* using available isolates from Latin America where previous studies are lacking. Our results indicate that the population structure of VR*Efm* in the region is diverse and can be grouped into two main lineages (Clades I and II) that belong to the previously reported clade A. A novel global reconstruction of the *E. faecium*, using a wide and diverse sample of isolates from 36 countries and obtained from clinical, animal, environmental and commensal samples, corroborates previous reports that recombination plays a major role in the evolution of this species. Our analyses also indicate, contrary to previous results, that animal-associated genomes are not monophyletic, and are instead a diverse collection of early-branching clades that diverged prior to the emergence of the human clinical clade and its two subclades (CRSI and CRSII).

The complex evolutionary dynamics of VR*Efm* highlight the importance of employing phylogenomic approaches when studying the population structure of a highly evolved hospital-associated pathogen.

## Acknowledgments

This work was founded by Universidad El Bosque, grant PCI 2016-8865 to L.D.; grants from the National Institutes of Health K24-AI121296 and R01-AI134637 to C.A.A. and grant FONDECYT regular Project No. 1171805, Ministry of Education, Government of Chile and the Millennium Science Initiative, Ministry of Economy, Development and Tourism, Chile to J.M.M.

## Methods

### *Enterococcus faecium* isolates

A total of 207 vancomycin-resistant Latin American *E. faecium* clinical isolates have been collected between 1998 and 2014 including those belonging to the first outbreak of VRE infections in Colombia and isolates collected in two multicentre surveillances^14,25,26^. Isolates were recovered from patients in Colombia (n=177, 86%) Peru (n=14, 7%), Venezuela (n=6, 3%), Ecuador (n=5, 2%) and Mexico (n=5, 2%). The most common sources included blood (22%), urine (18%) and stools (10%). For all the isolates, species (*E. faecium*) confirmation and the susceptibility profiles determination were performed by PCR assays^46^ and agar dilution, respectively^31^.

### Whole genome sequencing

From our VR*Efm* characterized strains collection, we selected 55 representative isolates based on distinct PFGE banding patterns. We included the first VRE reported in Colombia as the representative of an outbreak of 23 infections at a teaching hospital in 1998-1999^25^. Five isolates were selected from a national surveillance in Colombia during 2001-2002, which included 15 tertiary hospitals in 5 cities^26^ and 16 chosen from surveillance performed in Colombia, Ecuador, Venezuela and Peru in 2006-2008^14^. The remaining 33 isolates were sent to our lab for the confirmation of resistance or outbreak studies in 2005-2014. All selected isolates were recovered from clinical samples including blood (32%), urine (13%), faeces (13%), surgical wound (10%), pleural liquid (5%), peritoneal liquid (5%) and other sources (22%). The isolates were subjected to whole genome sequencing on the IIlumina platform. Briefly, genomic DNA was extracted from overnight cultures using the kit DNeasy Blood & Tissue Kit (Qiagen) after a lysozyme treatment. DNA libraries were prepared using Nextera XT kit (illumina) and sequenced on a MiSeq instrument using a 300pb paired-end strategy. The obtained paired-end reads were trimmed for quality and used for assemblies using SPAdes^47^.

### Global *E. faecium* genomic characterization

To place the population structure of Latin American VR*Efm* into global context, we included 285 *E. faecium* genomes from the publicly available collection available at NCBI. We aimed to incorporate a diverse set of sequences, including colonizing, commensal, animal and clinical sources recovered between 1946 and 2017 in Europe, North America, Asia, Africa, and Australia (Supplementary Table 1). Accordingly to the source, the *E. faecium* genomes were grouped into different categories: **i**) isolates from stools or rectal swabs of hospitalized patients (n=59), **ii**) organisms from hospitalized patients (n=196), recovered from sources other than faeces, including blood (n=113), urine (n=18) and other sources (n=65), **iii**) stools from healthy individuals not in hospital settings (n=13), **iv**) animal isolates (n=47), obtained from different animals, including pets, wild and farm animals, and **v**) “others” (n=25), which included isolates recovered from food products, water, soil, among other non-human and non-animal sources.

All sequences (340 *E. faecium* genomes) were annotated using RAST^48^. The sequence type (ST) was determined by MLST tools (https://github.com/tseemann/mlst) and verified against PubMLST^49^. Genomic characterization was performed to identify genetic elements associated with resistance using BLASTX^50^ searches against the ResFinder database^51^. Additionally, we specifically interrogated the genomes for substitutions in GyrAB and ParCE proteins associated to fluoroquinolone resistance, and mutations in genes encoding 23S rRNA and L3 and L4 proteins associated with linezolid resistance. Detection of mobile elements was performed with BlastN^50^. Search for *rep* families genes^52,53^ and insertion sequences (IS) was carried out with BLASTN searches and compared to the ISFinder database^54^. Identification of virulence elements was performed with BLASTX against a set of potential virulence proteins in enterococci^4,55^. Identification of CRISPR and *cas-*systems was done using CRISPRfinder^56^ and BLASTX searches using Cas system proteins^57^ as templates. All BLASTX hits were selected if they had an identity percentage higher or equal to 95% and a coverage of at least 80% of the target sequence. For BLASTN searches, hits were selected if they had an identity percentage higher than 90% and a coverage of at least 80% of the target sequence. To identify statistically significant differences across proportions of the evaluated characteristics among pairs of clades found, a Z-test was performed (α=0.01).

### Ampicillin resistance prediction based on PBP5 sequences

The ampicillin resistance prediction model for *E. faecium* isolates consisted on a random forest built upon a dataset of 250 PBP5 sequences from isolates with known MIC of ampicillin (62 from susceptible isolates [MIC≤8 µg/ml] and 188 belonging to resistant ones [MIC ≥ 16 µg/ml][Supplementary Table 4]). The model was based on a multiple sequence alignment using the sequence of the PBP5 from Com15 (GenBank accession: WP_002314979.1) isolate as reference (based on previous studies of correlation of the amino acid sequence of this protein with the resistant phenotype^34,35^) with 110 positions harbouring amino acid changes (Supplementary Table 4). These positions were used to create a random forest model with 100 decision trees; using a training set of 42 isolates (17 susceptible and 25 resistant with a range of MIC values). Based on this training set, forty amino acid changes were selected for the classification based on their discrimination power using recursive elimination process of those with lower score. Next, the model was tested on the whole dataset of PBP5 sequences and had a 100% specificity with 96% sensitivity, which resulted in 6 cases of major errors were the isolate was resistant but predicted to be susceptible.

### Phylogenetic analysis

We built a phylogenetic tree based on the core genome of 55 representative genomes from our collection, including the genome Com15 as outgroup. The core genome was obtained with Roary^58^ and each of the orthogroups was aligned with MUSCLE v3.8^59^. A Maximum Likelihood (ML) guide tree was built with RAxML 8.2.11^60^ using a GTR+G model. Using Bayesian approach, we estimated a Maximum Clade Credibility (MCC) tree based on 20 million trees in BEASTv1.8^61^. We employed a constant population size, a GTR+G+I substitution model, default prior probability distributions, and a chain length of 100 million steps with a burn-in of 10 million and a 5000-step thinning and the ML as starting tree.

The phylogenetic tree for the whole population of *E. faecium* included all the genomes (n=340) and two outgroups (*Enterococcus durans* BDGP3 [GenBank accession: CP022930.1] and *Enterococcus hirae* ATCC 9790 [CP003504.1]). This tree was based on the core genome (genes present in at least 90% of the studied genomes) obtained with Roary, each orthogroup was individually aligned with MUSCLE and then concatenated to obtain a matrix. The alignment matrix was used for Bayesian phylogenetic reconstruction with BEAST. Model parameters were the same as above with a chain length of 300 million steps, a burn-in of 80 million steps, and a random starting tree.

The second phylogenetic reconstruction included the genomes grouped into the clade corresponding to the previously designed Clade A^3^. We realized pairwise comparisons of the assemblies with Mummer 3.23^62^ against the reference genome Aus0085 (CP006620.1). The identified variants and the reference sequence were used to create a multiple whole genome alignment and, with it, we built a guide tree with RAxML^60^ using the abovementioned parameters. This guide tree was used later to obtain the recombinant regions in the alignment with ClonalFrameML^33^ for each isolate. Those regions were further removed from the alignment and then used to produce a MCC tree with BEAST. The same run parameters as above were used with a 50-million step burn-in.

Finally, a strict molecular clock analysis was performed on clade A strains. We dated the tips on the isolates accordingly to the sampling year. The analysis was done with the non-recombinant regions of the whole genome alignment as matrix and the MCC from the second analysis, as a guide tree. The analysis had a 300 million length chain and a burn-in of 30 million to obtain ESS numbers above 200. All MCC trees were computed with a 0.3 posterior clade probability cut-off and mean heights. To estimate the evolution rates across subclades, further subgrouping of the isolates was performed and a similar molecular clock analysis without guide tree were performed for each group using 100 million chain length and 10% burn in. All BEAST runs were performed on the CIPRES Science gateway servers^63^.

## Data Availability

All genomic data is available at GenBank database, accession numbers for the sequenced genomes are listed in Supplementary Table 3. The datasets generated during and/or analysed during the current study are available from the corresponding author on reasonable request.

## Ethics declarations

We declare no ethical competing interest. In our study, we did not perform any experiments with animals or higher invertebrates, neither performed experiments on humans and/or the use of human tissue samples. Our data have been originated from bacteria, not linked to clinical information, collected in previous studies and following full ethical approvals. Also, additional genomic data that we included for the analysis are available on public repositories (NCBI and published articles).

## Author contributions

R.R. performed experiments, carried out all statistical analyses, analysed results and wrote draft of the manuscript, L.D. and C.A.A conceived the study, analysed the results, and drafted and reviewed the manuscript, J.R. and D.P. conceived the study, interpreted data and analysed the results, P.J.P and SO.K. conceived experiments and provided key experimental suggestions, B.E.M. T.T.T and J.M.M interpreted and analyse data and helped to write the manuscript, L.P.C., S.R, A.M.E., A.D. and A.N. performed experiments and analysed data. All authors contributed to improve the manuscript and gave approval of the final version prior to submission.

## Additional Information

C.A.A has received grants funded by Merck Pharmaceuticals, MeMed Diagnostics Ltd and Entasis Therapeutics. B.E.M has received grants funded by Cubist/Merck, Forest/Actavis and is consultant of Paratek and Cempra.

The other authors declare no competing interests.

