## Supplemental material for "Genomic epidemiology of vancomycin resistant *Enterococcus faecium* (VR*Efm*) in Latin America: Revisiting the global VRE population structure" (from another version of this preprint: version 1, 2019-11-14)

[*]Correspondence

Lorena Diaz, PhD

Molecular Genetics and Antimicrobial Resistance Unit, International Center for Microbial Genomics, Universidad El Bosque

Av Cra 9 131a-02, Bogotá, Colombia.

**Supplementary Table 1.** List of *E. faecium* genomes included in the analysis

| GENOME | SOURCE CATEGORY | SOURCE | COUNTRY | YEAR | ST | SUBCLADE | ACCESSION |
|---|---|---|---|---|---|---|---|
| **ICIS96** | Animal | *Equus caballus* | Russia | 2014 | 47 | Animal branches | NNSQ01 |
| **E1630** | Other | River water | Netherlands | 1981 | 69 | Animal branches | AHXK01 |
| **E3346** | Hospitalized patient | Human - Blood | Netherlands | 2002 | 69 | Animal branches | AHYA01 |
| **E0679** | Animal | Pig | Belgium | 1995 | 150 | Animal branches | AHWM01 |
| **E0680** | Animal | Pig | Germany | 1995 | 151 | Animal branches | AHWN01 |
| **9930238_2** | Animal | Pig feces | Denmark | 1999 | 133 | Animal branches | AITJ01 |
| **841V03** | Animal | Pig feces | Denmark | 2003 | 147 | Animal branches | AISU01 |
| **E0688** | Animal | Pig | Spain | 1995 | 5 | Animal branches | AHWO01 |
| **HF50104** | Animal | Pig | USA | 2008 | 5 | Animal branches | AITR01 |
| **E1552** | Hospitalized patient feces | Human feces | Netherlands | 2002 | 5 | Animal branches | AHWW01 |
| **1_EFCM_CA_NL** | Hospitalized patient feces | Human feces | Netherlands | 2012 | 6 | Animal branches | NXOY01 |
| **E8SV3** | Animal | Pig feces | Denmark | 1995 | 6 | Animal branches | AITM01 |
| **7430166_3** | Animal | Pig feces | Denmark | 2002 | 6 | Animal branches | AIST01 |
| **9730219_1** | Animal | Pig feces | Denmark | 1997 | 6 | Animal branches | AISY01 |

| | | | | | | | |
|---|---|---|---|---|---|---|---|
| **7330519_3** | Animal | Pig feces | Denmark | 2001 | 6 | Animal branches | AISQ01 |
| **9830091_5** | Animal | Pig feces | Denmark | 1998 | 6 | Animal branches | AITC01 |
| **HF50106** | Animal | Pig feces | USA | 2008 | 6 | Animal branches | AITT01 |
| **7230532_1** | Animal | Pig feces | Denmark | 2000 | pst-incomplete | Animal branches | AISN01 |
| **A17SV1** | Animal | Pig feces | Denmark | 1995 | 6 | Animal branches | AITL01 |
| **IQ110** | Other | Cheese | Argentina | 2011 | SLV-52 | Animal branches | LKPG01 |
| **1231501** | Hospitalized patient | Human - Blood | USA | 2005 | 52 | Animal branches | ACAY01 |
| **E1258** | Hospitalized patient | Human - Blood | Spain | 1998 | 127 | Animal branches | AHWT01 |
| **5209** | Other | Chicken carcass rinse | USA | 2003 | 54 | Animal branches | NSDA01 |
| **7C8_DIV0219** | Animal | Marmot feces | Afghanistan | 2015 | New ST | Animal branches | NGKZ01 |
| **F1213D01** | Animal | *Bos taurus* feces | Canada | 2004 | SLV-955 | Animal branches | MJDZ01 |
| **D** | Other | Cheese | Mexico | 2008 | 1046 | Animal branches | MEGX01 |
| **VAN335** | Animal | Chicken feces | Denmark | 2010 | 417 | Animal branches | AIUZ01 |
| **E2071** | Animal | Poultry | Denmark | 2001 | 27 | Animal branches | AHXT01 |
| **E1574** | Animal | Dog | Belgium | 1995 | 27 | Animal branches | AHWY01 |

| | | | | | | | |
|---|---|---|---|---|---|---|---|
| **E0120** | Hospitalized patient | Human - Ascites | Netherlands | 1995 | 27 | Animal branches | AHWI01 |
| **E1185** | Hospitalized patient | Human - Blood | France | 1998 | 26 | Animal branches | AHWS01 |
| **E1679** | Hospitalized patient | Human - Vascular catheter | Brazil | 1998 | 114 | Animal branches | ABSC01 |
| **10_96A** | Hospitalized patient | Human - Blood | Brazil | 1996 | 281 | Animal branches | AXOL01 |
| **E1904** | Hospitalized patient | Human - Urine | Netherlands | 2001 | 210 | Animal branches | AHXQ01 |
| **E2134** | Animal | Chicken | Netherlands | 2004 | 12 | Animal branches | AHXU01 |
| **LS170308** | Animal | Forest musk deer (*Moschus berezovskii*) | China | 2017 | 30 | Animal branches | chromosome: NZ_CP025077.1/ CP025077.1 |
| **FDAARGOS_374** | Hospitalized patient | Human - Skin swab | USA | 2015 | New ST | Animal branches | PDFH01 |
| **7527** | Other | Chicken carcass rinse | USA | 2003 | 269 | Animal branches | NSDB01 |
| **E0269** | Animal | Turkey feces | Netherlands | 1996 | 9 | Animal branches | AHWK01 |
| **E1620** | Hospitalized patient | Human - Blood | Netherlands | 1957 | 67 | Animal branches | AHXF01 |
| **E0045** | Animal | Chicken feces | United Kingdom | 1992 | 9 | Animal branches | AHWH01 |
| **E1575** | Animal | Chicken | Belgium | 1995 | 158 | Animal branches | AHWZ01 |
| **E0164** | Animal | Turkey feces | Netherlands | 1996 | 26 | Animal branches | AHWJ01 |
| **F9730129_1** | Animal | Chicken | Denmark | 1997 | 245 | Animal branches | AITN01 |

| | | | | | | | |
|---|---|---|---|---|---|---|---|
| **E4215** | Animal | Chicken | Sweden | 2004 | 310 | Animal branches | AHYE01 |
| **2D4_DIV0473** | Animal | Buzzard | Germany | 2015 | 195 | Animal branches | NGLQ01 |
| **615** | Other | Environmental - Playground slide | USA | 2001 | 82 | Animal branches | NSDE01 |
| **E1627** | Hospitalized patient feces | Human feces | Netherlands | 1979 | 66 | Animal branches | AHXJ01 |
| **97_6_S5** | Hospitalized patient feces | Human feces | Chile | 2015 | 822 | Animal branches | MRYG01 |
| **WEFA23** | Healthy Individual Feces | Human - Healthy infants | China | 2015 | 66 | Animal branches | chromosome: CP021885.1 |
| **E1634** | Other | River water | Netherlands | 1982 | 66 | Animal branches | AHXL01 |
| **VAN345** | Animal | Chicken feces | Denmark | 2010 | 38 | Animal branches | AIVB01 |
| **E4452** | Animal | Dog feces | Netherlands | 2008 | 266 | Animal branches | AEOU01 |
| **E1626** | Hospitalized patient | Human - Stomach fluid | Netherlands | 1965 | 92 | Animal branches | AHXI01 |
| **GL3** | Animal | Dromedary milk | Algeria | 2013 | 22 | Animal branches | LIHM01 |
| **64_3** | Healthy Individual Feces | Human feces | Germany | 1992 | 21 | Animal branches | chromosome: NZ_CP012522.1/ CP012522.1 |
| **L_3** | Other | Probiotic | Russia | 2010 | 619 | Animal branches | JRGX01 |
| **9B11_DIV0348** | Animal | Bird | Germany | 2015 | 22 | Animal branches | NGKT01 |
| **70_36_8** | Animal | Pig | Denmark | 2011 | 536 | Animal branches | JRUG01 |

| | | | | | | | |
|---|---|---|---|---|---|---|---|
| **E1622** | Animal | Mouse | Netherlands | 1959 | 104 | Animal branches | AHXG01 |
| **E1050** | Healthy Individual Feces | Non-hospitalized individual Feces | Netherlands | 1998 | 92 | Animal branches | AHWQ01 |
| **NCTC13169** | Hospitalized patient | Human - Blood | United Kingdom | c. 1982 | 71 | Animal branches | ERR375583 |
| **NCTC7380** | Hospitalized patient | Human - Blood | United Kingdom | ≤ 1946 | 32 | Animal branches | ERR375572 |
| **E1071** | Hospitalized patient feces | Human feces | Netherlands | 2000 | 32 | Animal branches | ABQI01 |
| **ATCC8459** | Other | Cheese | USA | 1972 | 860 | Animal branches | AJAJ01 |
| **CRL1879** | Other | Cheese | Argentina | 2011 | New ST | Animal branches | AOUK01 |
| **11B6_DIV0698** | Animal | Mollusc | USA | 2013 | DLV-304 | Animal branches | NGMC01 |
| **E1573** | Animal | Turkey feces | Netherlands | 1996 | 21 | Animal branches | AHWX01 |
| **E1576** | Animal | Ostrich caecum | South Africa | 2001 | 159 | Animal branches | AHXA01 |
| **E1623** | Hospitalized patient | Human - Pus | Netherlands | 1960 | 22 | Animal branches | AHXH01 |
| **NCTC7171** | Hospitalized patient | Human - Blood | United Kingdom | ≤ 1946 | 160 | Animal branches | ERR375571 |
| **LCT_EF128** | Hospitalized patient | Human - Bronchoalveolar lavage | China | 2011 | 160 | Animal branches | AJUP01 |
| **E1578** | Animal | Pig feces | Germany | 2001 | 160 | Animal branches | AHXB01 |
| **471** | Hospitalized patient | Human - Blood | Russia | 2016 | SLV-168 | | PDFM01 |

| | | | | | | | |
|---|---|---|---|---|---|---|---|
| **993** | Hospitalized patient | Human - Wound | Russia | 2015 | SLV-168 | | PDLZ01 |
| **E161** | Hospitalized patient | Human - Urine | China | 2013 | 414 | | JXZA01 |
| **E31** | Hospitalized patient | Human - Bile | China | 2013 | 572 | | JYCO01 |
| **SC4** | Other | Fodder | China | 2016 | 78 | | chromosome:CP025425.1 |
| **BSAC_EC2783** | Hospitalized patient | Human - Blood | United Kingdom | 2011 | 412 | Clinical subclade I | ERR374675 |
| **ERV144** | Hospitalized patient | Human - Blood | Peru | 2007 | 494 | Clinical subclade I | AMAK01 |
| **ERV180** | Hospitalized patient | Human Pleural liquid | Colombia | 2010 | 656 | Clinical subclade I | MJFF00000000 |
| **MRSN3418** | Hospitalized patient | Human - Wound | USA | 2011 | 412 | Clinical subclade I | LBIM01 |
| **LIM1759** | Hospitalized patient feces | Human feces | Brazil | 2014 | SLV-896 | Clinical subclade I | MVGJ01 |
| **LIM1590** | Hospitalized patient feces | Human feces | Brazil | 2013 | 963 | Clinical subclade I | MVGG01 |
| **LIM1546** | Hospitalized patient feces | Human - Rectal swap | Brazil | 2013 | 896 | Clinical subclade I | MVGO01 |
| **LIM1586** | Hospitalized patient feces | Human feces | Brazil | 2013 | 896 | Clinical subclade I | MVGF01 |
| **LIM918** | Hospitalized patient feces | Human feces | Brazil | 2012 | 896 | Clinical subclade I | MVGE01 |
| **LIM4247** | Hospitalized patient feces | Human feces | Brazil | 2014 | 896 | Clinical subclade I | MVGH01 |
| **LIM1530** | Hospitalized patient feces | Human - Rectal swap | Brazil | 2013 | 896 | Clinical subclade I | MVGD01 |
| **IHC2** | Hospitalized patient | Human - Blood | USA | 2013 | 664 | Clinical subclade I | MSHR01 |

| | | | | | | | |
|---|---|---|---|---|---|---|---|
| **FDAARGOS_326** | Hospitalized patient feces | Human - jejunal aspirate | USA | 2015 | 736 | Clinical subclade I | POVW01 |
| **E39** | Hospitalized patient | Human - Blood | USA | 2009 | 736 | Clinical subclade I | chromosome: NZ_CP011281.1/ CP011281.1 |
| **ISMMS_VRE_10** | Hospitalized patient | Human - Blood | USA | 2011 | 736 | Clinical subclade I | chromosome: NZ_CP012471.1/ CP012471.1 |
| **ER04120.3A** | Hospitalized patient | Human - Blood | USA | 2015 | 736 | Clinical subclade I | chromosome: CP023784.1 |
| **ER03933.3A** | Hospitalized patient | Human - Blood | USA | 2015 | 736 | Clinical subclade I | chromosome: CP023780.1 |
| **ISMMS_VRE_5** | Hospitalized patient | Human - Blood | USA | 2011 | 736 | Clinical subclade I | chromosome: NZ_CP012454.1/ CP012454.1 |
| **E240** | Hospitalized patient | Human - Uterus cancer | USA | 2013 | 736 | Clinical subclade I | chromosome: NZ_CP017792.1/ CP017792.1 |
| **909_EFCM** | Hospitalized patient | Human - Urine | USA | 2012 | 736 | Clinical subclade I | JUOK01 |
| **702_EFCM** | Hospitalized patient | Human - Urine | USA | 2012 | 736 | Clinical subclade I | JUWL01 |
| **514_EFCM** | Hospitalized patient | Human - Urine | USA | 2012 | 736 | Clinical subclade I | JVEC01 |
| **377_EFCM** | Hospitalized patient | Human - Sputum induced | USA | 2012 | 736 | Clinical subclade I | JVJM01 |
| **43_EFCM** | Hospitalized patient | Human - Blood | USA | 2012 | 736 | Clinical subclade I | JVHJ01 |
| **41_EFCM** | Hospitalized patient | Human - Blood | USA | 2012 | SLV-736 | Clinical subclade I | JVIG01 |
| **97_7_S6** | Hospitalized patient feces | Human feces | Chile | 2015 | New ST | Clinical subclade I | MRYH01 |

| | | | | | | | |
|---|---|---|---|---|---|---|---|
| **BSAC_EC2288** | Hospitalized patient | Human - Blood | United Kingdom | 2010 | 78 | Clinical subclade I | ERR374760 |
| **197806558** | Hospitalized patient feces | Human feces | Sweden | 2014 | 192 | Clinical subclade I | LNLA01 |
| **VRE_1502382** | Hospitalized patient feces | Human feces | Sweden | 2015 | pst-incom plete | Clinical subclade I | LNOT01 |
| **VRE_1400136** | Hospitalized patient feces | Human feces | Sweden | 2014 | 192 | Clinical subclade I | LNLL01 |
| **S_1402282** | Hospitalized patient feces | Human feces | Sweden | 2014 | 192 | Clinical subclade I | LNLD01 |
| **E13931** | Hospitalized patient feces | Human feces | Sweden | 2013 | 192 | Clinical subclade I | LNLB01 |
| **ERV223** | Hospitalized patient | Human - Blood | Colombia | 2011 | 17 | Clinical subclade I | MJFL00000000 |
| **ERV218** | Hospitalized patient feces | Human - Rectal swap | Colombia | 2011 | 17 | Clinical subclade I | MJFJ00000000 |
| **2014_VREF_114** | Hospitalized patient feces | Human - Rectal swap | South Korea | 2014 | pst-incom plete | Clinical subclade I | chromosome: NZ_CP019970.1/ CP019970.1 |
| **2014_VREF_268** | Hospitalized patient feces | Human - Rectal swap | South Korea | 2015 | pst-incom plete | Clinical subclade I | chromosome: NZ_CP019992.1/ CP019992.1 |
| **EF_DMG150050 1** | Hospitalized patient | Human - No Data | Australia | 2014 | pst-incom plete | Clinical subclade I | chromosome 1:LT603678.1 |
| **VREF001** | Hospitalized patient | Human - Cholecystitis | United Kingdom | 2017 | pst-incom plete | Clinical subclade I | PJZU01 |
| **VREF005** | Hospitalized patient | Human - Colon cancer | United Kingdom | 2017 | pst-incom plete | Clinical subclade I | PJZQ01 |

| | | | | | | | |
|---|---|---|---|---|---|---|---|
| **VREF004** | Hospitalized patient | Human - Colon cancer | United Kingdom | 2017 | pst-incomplete | Clinical subclade I | PJZR01 |
| **VREF002** | Hospitalized patient | Human - Cholecystitis | United Kingdom | 2017 | pst-incomplete | Clinical subclade I | PJZT01 |
| **ERV242** | Hospitalized patient | Human - Urine | Colombia | 2012 | 412 | Clinical subclade I | MJFO00000000 |
| **ERV237** | Hospitalized patient feces | Human feces | Colombia | 2012 | 412 | Clinical subclade I | MJFN00000000 |
| **K59_68** | Hospitalized patient | Human - Blood | Norway | 2008 | 203 | Clinical subclade I | NMZH01 |
| **BSAC_EC2041** | Hospitalized patient | Human - Blood | United Kingdom | 2009 | 203 | Clinical subclade I | ERR375174 |
| **ERV251** | Hospitalized patient | Human - Osteomyelitis | Colombia | 2013 | 412 | Clinical subclade I | MJFP00000000 |
| **U_1313438** | Hospitalized patient feces | Human feces | Sweden | 2013 | DLV-203 | Clinical subclade I | LNLE01 |
| **S_1001508** | Hospitalized patient feces | Human feces | Sweden | 2010 | SLV-192 | Clinical subclade I | LNLC01 |
| **VRE1044** | Hospitalized patient | Human - Blood | Sweden | 2007 | 192 | Clinical subclade I | JAAL01 |
| **VRE1261** | Hospitalized patient | Human - Blood | Sweden | 2008 | SLV-192 | Clinical subclade I | JAAM01 |
| **C309** | Hospitalized patient | Human - Peritoneal fluid | China | 2006 | 78 | Clinical subclade I | AJTW01 |
| **XH877** | Hospitalized patient feces | Human feces | China | 2012 | 203 | Clinical subclade I | LXWZ01 |
| **VRE16** | Hospitalized patient | Human - Endocarditis | Brazil | 2009 | 412 | Clinical subclade I | LSYW01 |

| | | | | | | | |
|---|---|---|---|---|---|---|---|
| **ISMMS_VRE_2** | Hospitalized patient | Human - Blood | USA | 2011 | 656 | Clinical subclade I | chromosome: NZ_CP012436.1/ CP012436.1 |
| **LIM559** | Hospitalized patient feces | Human feces | Brazil | 2008 | 412 | Clinical subclade I | MXAS01 |
| **AUS0085** | Hospitalized patient | Human - Blood | Australia | 2009 | 203 | Clinical subclade I | chromosome: NC_021994.1/ CP006620.1 |
| **E6012** | Hospitalized patient feces | Human - Rectal swap | Latvia | 2010 | 78 | Clinical subclade I | AHYK01 |
| **BSAC_EC1359** | Hospitalized patient | Human - Blood | United Kingdom | 2006 | 78 | Clinical subclade I | ERR374663 |
| **E6045** | Hospitalized patient feces | Human - Rectal swap | Portugal | 2010 | 78 | Clinical subclade I | AHYL01 |
| **4686** | Hospitalized patient | Human - Blood | Russia | 2009 | 192 | Clinical subclade I | MQRF01 |
| **24** | Hospitalized patient feces | Human feces | Russia | 2012 | 78 | Clinical subclade I | PDMA01 |
| **24_10** | Hospitalized patient | Human - Wound Infection | Russia | 2011 | 78 | Clinical subclade I | MVEB01 |
| **U0317** | Hospitalized patient | Human - Urinary tract infection | Netherlands | 2005 | 78 | Clinical subclade I | ABSW01 |
| **E2560** | Hospitalized patient | Human - Blood | Netherlands | 2006 | 78 | Clinical subclade I | AHYI01 |
| **ERV161** | Hospitalized patient | Human - Blood | Colombia | 2008 | 203 | Clinical subclade I | AMAW01 |
| **1231502** | Hospitalized patient | Human - Blood | USA | 2005 | 203 | Clinical subclade I | ACAX01 |
| **ERV108** | Hospitalized patient | Human - Urine | Ecuador | 2006 | 203 | Clinical subclade I | AMAZ01 |
| **ERV98** | Hospitalized patient | Human - Blood | Colombia | 2005 | 412 | Clinical subclade I | MJFY00000000 |

| | | | | | | | |
|---|---|---|---|---|---|---|---|
| **ERV89** | Hospitalized patient | Human - Bronchial secretion | Colombia | 2005 | 412 | Clinical subclade I | MTRT00000000 |
| **ERV114** | Hospitalized patient | Human - Peritoneal liquid | Colombia | 2006 | 412 | Clinical subclade I | AMBB01 |
| **ERV102** | Hospitalized patient | Human - Sputum | Colombia | 2006 | 412 | Clinical subclade I | AMAX01 |
| **ERV168** | Hospitalized patient | Human - Surgical wound secretion | Colombia | 2009 | 412 | Clinical subclade I | AMAU01 |
| **ERV172** | Hospitalized patient | Human - No data | Colombia | 2009 | 412 | Clinical subclade I | MJFB00000000 |
| **ERV220** | Hospitalized patient | Human - No data | Colombia | 2011 | 412 | Clinical subclade I | MJFK00000000 |
| **ERV174** | Hospitalized patient | Human - Peritoneal liquid | Colombia | 2009 | 412 | Clinical subclade I | MJFC00000000 |
| **ERV177** | Hospitalized patient feces | Human - Rectal swap | Colombia | 2010 | 412 | Clinical subclade I | MJFE00000000 |
| **805447_07** | Hospitalized patient | Human - Pleura effusion | Austria | 2007 | 412 | Clinical subclade I | LQRS01 |
| **IHC11** | Hospitalized patient | Human - Blood | USA | 2007 | 584 | Clinical subclade I | MSIB01 |
| **10_131** | Hospitalized patient | Human - Catheter | France | 2010 | 412 | Clinical subclade I | PHKY01 |
| **967_EFCM** | Hospitalized patient | Human - Wound | USA | 2012 | 412 | Clinical subclade I | JULR01 |
| **964_EFCM** | Hospitalized patient | Human - Fluid | USA | 2012 | 412 | Clinical subclade I | JULU01 |
| **990_EFCM** | Hospitalized patient | Human - Fluid | USA | 2012 | 412 | Clinical subclade I | JUKT01 |
| **LIM716** | Hospitalized patient feces | Human feces | Brazil | 2008 | DLV-478 | Clinical subclade I | MVGL01 |

| | | | | | | | |
|---|---|---|---|---|---|---|---|
| **LIM1547** | Hospitalized patient feces | Human - Rectal swap | Brazil | 2013 | 478 | Clinical subclade I | MVGP01 |
| **LIM554** | Hospitalized patient | Human - Blood | Brazil | 2008 | 412 | Clinical subclade I | MVGM01 |
| **LIM563** | Hospitalized patient feces | Human feces | Brazil | 2009 | 412 | Clinical subclade I | MVGK01 |
| **LIM699** | Hospitalized patient | Human - Blood | Brazil | 2006 | 412 | Clinical subclade I | MXAU01 |
| **LIM695** | Hospitalized patient | Human - Blood | Brazil | 2006 | 478 | Clinical subclade I | MXAT01 |
| **ERV124** | Hospitalized patient | Human - Surgical wound secretion | Peru | 2007 | 412 | Clinical subclade I | AMAM01 |
| **IHC8** | Hospitalized patient | Human - Blood | USA | 2007 | 412 | Clinical subclade I | MSHY01 |
| **IHC13** | Hospitalized patient | Human - Blood | USA | 2006 | 412 | Clinical subclade I | MSID01 |
| **ERV133** | Hospitalized patient | Human - Urine | Venezuela | 2007 | 412 | Clinical subclade I | MJEY00000000 |
| **ERV118** | Hospitalized patient | Human - Blood | Venezuela | 2006 | 412 | Clinical subclade I | MTRU00000000 |
| **ERV115** | Hospitalized patient | Human - Surgical wound secretion | Venezuela | 2006 | 412 | Clinical subclade I | MTRV00000000 |
| **463_ANA** | Hospitalized patient | Human - Blood | USA | 2015 | SLV-750 | Clinical subclade I | NETE01 |
| **463_AER** | Hospitalized patient | Human - Blood | USA | 2015 | 750 | Clinical subclade I | NETD01 |
| **ERV175** | Hospitalized patient | Human - Urine | Colombia | 2009 | 412 | Clinical subclade I | MJFD00000000 |
| **ERV155** | Hospitalized patient | Human - Blood | Colombia | 2008 | 412 | Clinical subclade I | MJEZ00000000 |
| **ERV165** | Hospitalized patient feces | Human feces | Colombia | 2008 | 412 | Clinical subclade I | AMAV01 |

| | | | | | | | |
|---|---|---|---|---|---|---|---|
| **ERV157** | Hospitalized patient feces | Human feces | Colombia | 2008 | 412 | Clinical subclade I | MJFA00000000 |
| **ERV196** | Hospitalized patient | Human - Blood | Colombia | 2010 | 412 | Clinical subclade I | MJFH00000000 |
| **VRE_1502939** | Hospitalized patient feces | Human feces | Sweden | 2015 | New ST | Clinical subclade II | LNOW01 |
| **VRE2014_7** | Hospitalized patient feces | Human - Rectal swap | USA | 2014 | New ST | Clinical subclade II | NLCP01 |
| **VREN1625** | Hospitalized patient | Human - Blood | United Kingdom | 2015 | 80 | Clinical subclade II | FXHD01 |
| **VREN1371** | Hospitalized patient | Human - Blood | United Kingdom | 2015 | 80 | Clinical subclade II | FXHP01 |
| **ICBEF1B** | Other | Seafood | Brazil | 2016 | 1336 | Clinical subclade II | NKYP01 |
| **LIM4275** | Hospitalized patient feces | Human feces | Brazil | 2014 | 792 | Clinical subclade II | MVGI01 |
| **ET13_17** | Hospitalized patient feces | Human - Rectal swap | Brazil | 2017 | New ST | Clinical subclade II | PEJX02 |
| **ET12_17** | Hospitalized patient | Human - Blood | Brazil | 2017 | New ST | Clinical subclade II | PEJV02 |
| **ET14_17** | Hospitalized patient feces | Human - Rectal swap | Brazil | 2017 | New ST | Clinical subclade II | PEJY02 |
| **ERV190** | Hospitalized patient | Human - No data | Colombia | 2010 | 1517 | Clinical subclade II | MJFG00000000 |
| **BSAC_EC1822** | Hospitalized patient | Human - Blood | United Kingdom | 2008 | 412 | Clinical subclade II | ERR374884 |
| **BSAC_EC2499** | Hospitalized patient | Human - Blood | United Kingdom | 2010 | 412 | Clinical subclade II | ERR374686 |
| **FDAARGOS_332** | Hospitalized patient feces | Human - Stool sample | USA | 2015 | 17 | Clinical subclade II | NJFI02 |
| **VRE2014_195** | Hospitalized patient feces | Human - Rectal swap | USA | 2014 | 17 | Clinical subclade II | NLCQ01 |

| | | | | | | | |
|---|---|---|---|---|---|---|---|
| **VRE2016_194** | Hospitalized patient feces | Human - Rectal swap | USA | 2016 | 17 | Clinical subclade II | NLCT01 |
| **VRE001** | Hospitalized patient | Human - Blood | USA | 2013 | 203 | Clinical subclade II | chromosome: NZ_CP018071.1/ CP018071.1 |
| **712_EFCM** | Hospitalized patient | Human - Urine | USA | 2012 | 412 | Clinical subclade II | JUWB01 |
| **969_EFCM** | Hospitalized patient | Human - Urine | USA | 2012 | 412 | Clinical subclade II | JULP01 |
| **BSAC_EC2198** | Hospitalized patient | Human - Blood | United Kingdom | 2009 | 665 | Clinical subclade II | ERR374807 |
| **BSAC_EC1484** | Hospitalized patient | Human - Blood | United Kingdom | 2006 | 665 | Clinical subclade II | ERR374754 |
| **1230933** | Hospitalized patient | Human - Blood | USA | 2005 | 18 | Clinical subclade II | ACAS01 |
| **ISMMS_VRE_11** | Hospitalized patient | Human - Blood | USA | 2011 | 18 | Clinical subclade II | chromosome: NZ_CP016163.1 CP016163.1 |
| **VRE2015_149** | Hospitalized patient feces | Human - Rectal swap | USA | 2015 | 18 | Clinical subclade II | NLCR01 |
| **FDAARGOS_138** | Hospitalized patient feces | Human - Stool sample | USA | 2014 | 18 | Clinical subclade II | LORO02 |
| **MRSN11639** | Hospitalized patient | Human - Wound | USA | 2012 | 18 | Clinical subclade II | LBIN01 |
| **199_EFCM** | Hospitalized patient | Human - Blood | USA | 2012 | 18 | Clinical subclade II | JVQH01 |
| **999_EFCM** | Hospitalized patient | Human - Abdominal Fluid | USA | 2012 | 18 | Clinical subclade II | JUKJ01 |
| **715_EFCM** | Hospitalized patient | Human - Urine | USA | 2012 | 18 | Clinical subclade II | JUVY01 |
| **704_EFCM** | Hospitalized patient | Human - Blood | USA | 2012 | 18 | Clinical subclade II | JUWJ01 |

| | | | | | | | |
|---|---|---|---|---|---|---|---|
| **VRE_1502913** | Hospitalized patient feces | Human feces | Sweden | 2015 | 117 | Clinical subclade II | LNOV01 |
| **E2** | Hospitalized patient | Human - Urine | Spain | 2011 | 117 | Clinical subclade II | MPZY01 |
| **1250_EFCM** | Hospitalized patient | Human - Blood | USA | 2012 | 117 | Clinical subclade II | JVWZ01 |
| **1232_EFCM** | Hospitalized patient | Human - Blood | USA | 2012 | 117 | Clinical subclade II | JVXT01 |
| **1238_EFCM** | Hospitalized patient | Human - Blood | USA | 2012 | 117 | Clinical subclade II | JVXN01 |
| **97_3_S3** | Hospitalized patient feces | Human feces | Chile | 2015 | New ST | Clinical subclade II | MRYF01 |
| **EF_AUS00233** | Healthy Individual Feces | Human feces | Australia | 2011 | 796 | Clinical subclade II | chromosome 1: NZ_LT598663.1/ LT598663.1 |
| **18434** | Hospitalized patient | Human - Blood | Russia | 2012 | 78 | Clinical subclade II | PDFL01 |
| **ERS1122499** | Hospitalized patient | Human - Blood | Germany | 2013 | 17 | Clinical subclade II | FKLC01 |
| **9_F_6** | Hospitalized patient | Human - Blood | Norway | 2012 | 872 | Clinical subclade II | NMZI01 |
| **MRSN4777** | Hospitalized patient | Human - Wound | USA | 2011 | No typeable | Clinical subclade II | LBIL01 |
| **560_EFCM** | Hospitalized patient | Human - Blood | USA | 2012 | SLV-923 | Clinical subclade II | JVCC01 |
| **VRE1558** | Hospitalized patient | Human - Blood | Brazil | 2013 | No typeable | Clinical subclade II | LBCV01 |
| **ERV280** | Hospitalized patient | Human - Urine | Colombia | 2014 | 1516 | Clinical subclade II | MJFU00000000 |

| | | | | | | | |
|---|---|---|---|---|---|---|---|
| **BSAC_EC1116** | Hospitalized patient | Human - Blood | United Kingdom | 2005 | 202 | Clinical subclade II | ERR374680 |
| **BSAC_EC2579** | Hospitalized patient | Human - Blood | United Kingdom | 2011 | 117 | Clinical subclade II | ERR374756 |
| **HPH2** | Hospitalized patient | Human - Urinary tract infection | Portugal | 2007 | 125 | Clinical subclade II | MBRH01 |
| **E1731** | Hospitalized patient | Human - Blood | Tanzania | 2002 | 18 | Clinical subclade II | AHXO01 |
| **ERV275** | Hospitalized patient | Human - No data | Mexico | 2012 | 1304 | Clinical subclade II | MJFS00000000 |
| **BSAC_EC1179** | Hospitalized patient | Human - Blood | United Kingdom | 2005 | 1042 | Clinical subclade II | ERR374732 |
| **BSAC_EC1142** | Hospitalized patient | Human - Blood | United Kingdom | 2005 | 64 | Clinical subclade II | ERR374989 |
| **BSAC_EC1201** | Hospitalized patient | Human - Blood | United Kingdom | 2005 | pst-incomplete | Clinical subclade II | ERR374823 |
| **BSAC_EC956** | Hospitalized patient | Human - Blood | United Kingdom | 2004 | pst-incomplete | Clinical subclade II | ERR375172 |
| **76** | Hospitalized patient | Human - Skin | Russia | 2014 | 80 | Clinical subclade II | PDNS01 |
| **97_19_S17** | Hospitalized patient feces | Human feces | Chile | 2015 | 262 | Clinical subclade II | MRYE01 |
| **ERV279** | Hospitalized patient | Human - No data | Mexico | 2012 | 770 | Clinical subclade II | MJFT00000000 |
| **2E6_DIV0576** | Animal | Bird | Germany | 2015 | 80 | Clinical subclade II | NGLP01 |
| **93** | Hospitalized patient | Human - Skin | Russia | 2013 | 18 | Clinical subclade II | PDFN01 |
| **BSAC_EC1514** | Hospitalized patient | Human - Blood | United Kingdom | 2007 | 18 | Clinical subclade II | ERR374938 |

| | | | | | | | |
|---|---|---|---|---|---|---|---|
| **32** | Hospitalized patient | Human - Burn wound | Russia | 2010 | 18 | Clinical subclade II | PDFP01 |
| **ST_18_K073** | Hospitalized patient | Human - Urine | South Africa | 2013 | 18 | Clinical subclade II | NXIX01 |
| **607_EFCM** | Hospitalized patient | Human - Fluid | USA | 2012 | 18 | Clinical subclade II | JVAH01 |
| **E2883** | Hospitalized patient | Human - Blood | Netherlands | 2002 | 18 | Clinical subclade II | AHXX01 |
| **BSAC_EC824** | Hospitalized patient | Human - Blood | United Kingdom | 2004 | 18 | Clinical subclade II | ERR374662 |
| **BSAC_EC1651** | Hospitalized patient | Human - Blood | United Kingdom | 2007 | 78 | Clinical subclade II | ERR374849 |
| **ERV122** | Hospitalized patient | Human - Surgical wound secretion | Peru | 2007 | 18 | Clinical subclade II | AMAO01 |
| **ERV99** | Hospitalized patient | Human - Peritoneal liquid | Colombia | 2005 | 18 | Clinical subclade II | AMAQ01 |
| **ERV138** | Hospitalized patient | Human - Blood | Colombia | 2007 | 18 | Clinical subclade II | AMBD01 |
| **ERV109** | Hospitalized patient | Human - Blood | Colombia | 2006 | 18 | Clinical subclade II | AMBC01 |
| **E1321** | Hospitalized patient | Human - Catheter | Italy | 1999 | 78 | Clinical subclade II | AHYF01 |
| **E4389** | Animal | Dog feces | Denmark | 2008 | 78 | Clinical subclade II | AHYJ01 |
| **E4453** | Animal | Dog feces | Netherlands | 2008 | 192 | Clinical subclade II | AEDZ01 |
| **FDAARGOS_331** | Hospitalized patient feces | Human - Stool sample | USA | 2015 | 18 | Clinical subclade II | POVX01 |
| **BSAC_EC832** | Hospitalized patient | Human - Blood | United Kingdom | 2004 | 186 | Clinical subclade II | ERR375187 |
| **E1133** | Hospitalized patient feces | Human feces | USA | 2001 | 117 | Clinical subclade II | AHWR01 |

| | | | | | | | |
|---|---|---|---|---|---|---|---|
| **1231408** | Hospitalized patient | Human - Blood | USA | 2005 | 582 | Clinical subclade II | ACBB01 |
| **E0333** | Hospitalized patient | Human - Blood | Israel | 1997 | 80 | Clinical subclade II | AHWL01 |
| **VRE2** | Hospitalized patient | Human - Blood | Malaysia | 2011 | 80 | Clinical subclade II | LTAA01 |
| **VRER5** | Hospitalized patient | Human - Endocarditis | Malaysia | 2011 | 80 | Clinical subclade II | LTBJ01 |
| **TX16_DO** | Hospitalized patient | Human - Blood | USA | 1992 | 18 | Clinical subclade II | chromosome: NC_017960.1/ CP003583.1 |
| **E1392** | Hospitalized patient feces | Human feces | United Kingdom | 2000 | 64 | Clinical subclade II | AHWV01 |
| **NCTC12202** | Hospitalized patient | Human - Blood | United Kingdom | 1986 | 18 | Clinical subclade II | ERR375579 |
| **NCTC12204** | Hospitalized patient | Human - Blood | United Kingdom | 1986 | 18 | Clinical subclade II | ERR375585 |
| **ISMMS_VRE_8** | Hospitalized patient | Human - Blood | USA | 2011 | 17 | Clinical subclade II | chromosome: NZ_CP012465.1/ CP012465.1 |
| **ERV281** | Hospitalized patient | Human - Blood | Colombia | 2014 | 17 | Clinical subclade II | MJFV00000000 |
| **ERV231** | Hospitalized patient | Human - Blood | Colombia | 2012 | 17 | Clinical subclade II | MJFM00000000 |
| **ERV212** | Hospitalized patient | Human - Blood | Colombia | 2011 | 17 | Clinical subclade II | MTRX00000000 |
| **ERV214** | Hospitalized patient feces | Human feces | Colombia | 2011 | 17 | Clinical subclade II | MJFI00000000 |
| **ERV263** | Hospitalized patient | Human - Blood | Colombia | 2013 | 17 | Clinical subclade II | MJFQ00000000 |
| **ERV270** | Hospitalized patient feces | Human feces | Colombia | 2013 | 17 | Clinical subclade II | MJFR00000000 |

| | | | | | | | |
|---|---|---|---|---|---|---|---|
| **ERV28** | Hospitalized patient | Human - Surgical wound secretion | Colombia | 2001 | 17 | Clinical subclade II | MTRY00000000 |
| **ERV35** | Hospitalized patient | Human - Blood | Colombia | 2001 | 17 | Clinical subclade II | MJFX00000000 |
| **ERV30** | Hospitalized patient | Human - Surgical wound secretion | Colombia | 2001 | 17 | Clinical subclade II | MTRW00000000 |
| **ERV26** | Hospitalized patient | Human - Pleural liquid | Colombia | 2001 | 17 | Clinical subclade II | AMAT01 |
| **ERV34** | Hospitalized patient | Human - No data | Colombia | 2001 | 17 | Clinical subclade II | MJFW00000000 |
| **ERV69** | Hospitalized patient | Human - Fistula | Colombia | 2005 | 17 | Clinical subclade II | AMAR01 |
| **ERV38** | Hospitalized patient | Human - No data | Colombia | 2002 | 17 | Clinical subclade II | AMAS01 |
| **TX0082** | Hospitalized patient | Human - Blood | USA | 1999 | 17 | Clinical subclade II | AEBU01 |
| **LIM714** | Hospitalized patient feces | Human feces | Brazil | 2008 | New ST | Clinical subclade II | MVGN01 |
| **E1644** | Hospitalized patient | Human - Catheter-Urine | Germany | 2002 | 78 | Clinical subclade II | AHYG01 |
| **E2369** | Hospitalized patient | Human - Wound | Hungary | 2005 | 78 | Clinical subclade II | AHYH01 |
| **E6** | Hospitalized patient feces | Human feces | Spain | 2009 | 17 | Clinical subclade II | MQAC01 |
| **AUS0004** | Hospitalized patient | Human - Blood | Australia | 1998 | 17 | Clinical subclade II | chromosome: NC_017022.1/ CP003351.1 |
| **E525** | Hospitalized patient | Human - Wound | Australia | 1998 | 17 | Clinical subclade II | AUWV01 |
| **ERV121** | Hospitalized patient | Human - Blood | Peru | 2007 | 280 | Clinical subclade II | AMAP01 |

| | | | | | | | |
|---|---|---|---|---|---|---|---|
| **ERV123** | Hospitalized patient | Human - Urine | Peru | 2007 | 280 | Clinical subclade II | AMAN01 |
| **ERV125** | Hospitalized patient | Human - Blood | Peru | 2007 | 125 | Clinical subclade II | AMAL01 |
| **ERV113** | Hospitalized patient | Human - Urine | Peru | 2006 | 280 | Clinical subclade II | MJEW00000000 |
| **E155** | Healthy Individual Feces | Human - Stool sample | USA | 1995 | 17 | Clinical subclade II | AUWX01 |
| **BSAC_EC239** | Hospitalized patient | Human - Blood | United Kingdom | 2001 | 17 | Clinical subclade II | ERR374681 |
| **BSAC_EC227** | Hospitalized patient | Human - Blood | United Kingdom | 2001 | 17 | Clinical subclade II | ERR374688 |
| **BSAC_EC754** | Hospitalized patient | Human - Blood | United Kingdom | 2004 | 17 | Clinical subclade II | ERR374714 |
| **VRE0576** | Hospitalized patient | Human - Blood | Sweden | 2006 | 17 | Clinical subclade II | JAAK01 |
| **E1162** | Hospitalized patient | Human - Blood | France | 1997 | 17 | Clinical subclade II | ABQJ01 |
| **ATCC51559** | Hospitalized patient | Human - Blood | USA | 1992 | 17 | Clinical subclade II | JSVT01 |
| **E1165** | Hospitalized patient | Human - Wound | Italy | 1997 | 17 | Clinical subclade II | AUWW01 |
| **1231410** | Hospitalized patient | Human - Skin and soft tissue | USA | 2005 | 17 | Clinical subclade II | ACBA01 |
| **E2297** | Hospitalized patient | Human - Urine | USA | 2001 | 117 | Clinical subclade II | AHXV01 |
| **ERV1** | Hospitalized patient | Human - Pleural liquid | Colombia | 1998 | 17 | Clinical subclade II | AMAY01 |
| **ERV107** | Hospitalized patient | Human - Blood | Ecuador | 2006 | 17 | Clinical subclade II | AMBA01 |

| | | | | | | | |
|---|---|---|---|---|---|---|---|
| **ERV127** | Hospitalized patient | Human - Blood | Ecuador | 2007 | 17 | Clinical subclade II | MJEX00000000 |
| **VRE3** | Hospitalized patient | Human - Blood | USA | 1998 | SLV-16 | Clinical subclade II | JSET01 |
| **C68** | Hospitalized patient feces | Human feces | USA | 1997 | 16 | Clinical subclade II | LPUE01 |
| **BSAC_EC426** | Hospitalized patient | Human - Blood | United Kingdom | 2002 | 16 | Clinical subclade II | ERR374669 |
| **BSAC_EC495** | Hospitalized patient | Human - Blood | United Kingdom | 2002 | 16 | Clinical subclade II | ERR374844 |
| **1141733** | Hospitalized patient | Human - Wound | USA | 2005 | 327 | Clade B | ACAZ01 |
| **17OM39** | Healthy Individual Feces | Human - Stool sample | India | 2015 | 76 | Clade B | LWHF01 |
| **7H8_DIV0219** | Animal | Marmot feces | Afghanistan | 2015 | New ST | Clade B | NGKW01 |
| **825** | Animal | Dairy cattle feces | USA | 2004 | SLV-94 | Clade B | NSDF01 |
| **BM4107** | Hospitalized patient feces | Human feces | France | 1988 | 172 | Clade B | JMII01 |
| **BSAC_EC555** | Hospitalized patient | Human - Blood | United Kingdom | 2003 | 1036 | Clade B | ERR374785 |
| **COM12** | Healthy Individual Feces | Non-hospitalized individual Feces | USA | 2006 | 107 | Clade B | ACBC01 |
| **COM15** | Healthy Individual Feces | Non-hospitalized individual Feces | USA | 2007 | 583 | Clade B | ACBD01 |
| **E1007** | Healthy Individual Feces | Non-hospitalized individual Feces | Netherlands | 1998 | 61 | Clade B | AHWP01 |

| | | | | | | | |
|---|---|---|---|---|---|---|---|
| **E1590** | Healthy Individual Feces | Non-hospitalized individual Feces | Ireland | 2001 | 163 | Clade B | AHXC01 |
| **E1604** | Other | Cheese | Norway | 1956 | 75 | Clade B | AHXD01 |
| **E1613** | Other | Fish burger | Norway | 1964 | 77 | Clade B | AHXE01 |
| **E1861** | Hospitalized patient feces | Human feces | Spain | 2001 | 289 | Clade B | AHXP01 |
| **E1972** | Hospitalized patient | Human - Blood | Germany | 2000 | 94 | Clade B | AHXR01 |
| **E2039** | Hospitalized patient | Human - K-spitze | Germany | 2000 | 296 | Clade B | AHXS01 |
| **E2620** | Hospitalized patient | Human - Blood | Netherlands | 2006 | 1175 | Clade B | AHXW01 |
| **E3083** | Hospitalized patient | Human - Blood | Netherlands | 2000 | 327 | Clade B | AHXZ01 |
| **E3548** | Hospitalized patient | Human - Blood | Netherlands | 2004 | 328 | Clade B | AHYB01 |
| **E980** | Healthy Individual Feces | Human - Healthy volunteer | Netherlands | 1998 | 94 | Clade B | ABQA01 |
| **F1129D110** | Animal | Bos taurus feces | Canada | 2004 | New ST | Clade B | MJDY01 |
| **GM75** | Other | Cheese | Argentina | 2011 | 94 | Clade B | LKPI01 |
| **HP_5_10** | Healthy Individual Feces | Human feces | Russia | 2016 | New ST | Clade B | PCGB01 |
| **JB00008** | Other | Korean fermented soybean paste (cheonggukjang) | South Korea | 2015 | 94 | Clade B | NPOO01 |
| **KACC15689** | Other | Doenjang fermented soybean paste | South Korea | 2013 | New ST | Clade B | LBIT01 |

| | | | | | | | |
|---|---|---|---|---|---|---|---|
| **KACC15700** | Other | Ganjang fermented soy sauce | South Korea | 2013 | New ST | Clade B | LDNC01 |
| **KACC15711** | Other | Meju fermented soybeans | South Korea | 2013 | New ST | Clade B | LDND01 |
| **KACC15960** | Other | Doenjang fermented soybean paste | South Korea | 2013 | New ST | Clade B | LDNE01 |
| **KACC16093** | Other | Meju fermented soybeans | South Korea | 2013 | New ST | Clade B | LDNH01 |
| **L_X** | Other | Probiotic preparation | Russia | 2010 | 812 | Clade B | JRGY01 |
| **M3K31** | Animal | Griffon vulture | Spain | 2010 | 76 | Clade B | LAXK01 |
| **NCTC7378** | Hospitalized patient | Human - Blood | United Kingdom | ≤ 1946 | 361 | Clade B | ERR375567 |
| **PC4_1** | Healthy Individual Feces | Non-hospitalized individual Feces | Australia | 2008 | 720 | Clade B | ADMM01 |
| **TX1330** | Healthy Individual Feces | Non-hospitalized individual Feces | USA | 2006 | 107 | Clade B | ACHL01 |
| **11F9MSG5001** | Animal | Chelonia mydas feces | Brazil | 0 | 688 | Clade B | NGMK01 |
| **UBA4566** | Other | Wood | USA | 2015 | 946 | Clade B | DGKQ01 |
| **UC7256** | Other | Food | Italy | 0 | 74 | Clade B | AWWM01 |
| **UC8668** | Other | Food | Italy | 0 | New ST | Clade B | AWWN01 |

**Supplementary table 2.** Genomic characteristics of the clades A and B of *E. faecium* population.

| | **CLADE A** | **CLADE B** |
|---|---|---|
| **Genome size** | 2.908 Mb ± 0.19 Mb | 2.833 Mb ± 0.16 Mb |
| **Gene content** | 2801 ± 208 genes | 2736 ± 156 genes |
| **Pan genome** | 10382 orthogroups | 23746 orthogroups |
| **Core genome** | 1182 orthogroups | 1466 orthogroups |
| **Average number of virulence genes** | 21.6 | 13.7 |
| **Average number of *rep* plasmids genes** | 3.24 | 2.16 |
| **Average number of insertion sequences (is)** | 7.4 | 4.1 |

**Supplementary table 3.** Comparison of the prevalences and frequencies of genetic elements related to antibiotic resistance, virulence, mobile elements, plasmid families and CRISPR/CAS-system among the genomes from the subgroups in the clade A.

| Category | Description | Element | Animal branches (n=73) | CRS-I (n=101) | CRS-II (n=124) | P value of the difference of proportions between | |
|---|---|---|---|---|---|---|---|
| | | | % (n) | % (n) | % (n) | Animal vs CRS-I | Animal vs CRS-II |
| Resistance | Glycopeptides resistance | VanA | 32,8 (24) | 70,2 (71) | 65,3 (81) | < .001 | < .001 |
| | | VanB | 0 (0) | 11,8 (12) | 9,6 (12) | < .001 | < .001 |
| | | VanD | 1,3 (1) | 0 (0) | 0 (0) | .246 | .246 |
| | | Total | 34,2 (25) | 82,1 (83) | 75 (93) | < .001 | < .001 |
| | Aminoglycosides resistance | aac(6')-aph(2'') | 6,8 (5) | 52,4 (53) | 47,5 (59) | < .001 | < .001 |
| | | ant(6)-Ia | 16,4 (12) | 74,2 (75) | 71,7 (89) | < .001 | < .001 |
| | | aph(3')-III | 12,3 (9) | 82,1 (83) | 71,7 (89) | < .001 | < .001 |
| | | aadD, aph(2'')-Ib, aph(2'')-Id, spc or str | 15 (11) | 7,9 (8) | 4 (5) | .143 | .021 |
| | | Total | 21,9 (16) | 90 (91) | 81,4 (101) | < .001 | < .001 |
| | Fluroquinolones resistance | GyrA mutations | 1,3 (1) | 96 (97) | 86,2 (107) | < .001 | < .001 |
| | | ParC mutations | 2,7 (2) | 100 (101) | 86,2 (107) | < .001 | < .001 |
| | | Total | 2,7 (2) | 100 (101) | 98,3 (122) | < .001 | < .001 |
| | Ampicillin resistance | PBP5-R | 9,5 (7) | 94 (95) | 96,7 (120) | < .001 | < .001 |
| | Trimethoprim resistance | dfrG | 4,1 (3) | 13,8 (14) | 53,2 (66) | .025 | < .001 |
| | MLSb resistance | erm(A) | 5,4 (4) | 6,9 (7) | 3,2 (4) | .366 | .309 |
| | | erm(B) | 39,7 (29) | 85,1 (86) | 83 (103) | < .001 | < .001 |
| | | erm(G) | 0 (0) | 0 (0) | 0,8 (1) | 1 | .241 |
| | | erm(T) | 1,3 (1) | 0,9 (1) | 10,4 (13) | .387 | .004 |
| | | lnu(B) | 9,5 (7) | 25,7 (26) | 4,8 (6) | .005 | .195 |

| | | | | | | | | |
|---|---|---|---|---|---|---|---|---|
| | | msr(C) | | 95,8 (70) | 100 (101) | 97,5 (121) | .08 | .328 |
| | | vat(D) | | 6,8 (5) | 0 (0) | 0 (0) | .027 | .027 |
| | | vat(E) | | 5,4 (4) | 0 (0) | 0 (0) | .049 | .049 |
| | | Total | | 98,6 (72) | 100 (101) | 100 (124) | .237 | .237 |
| | Tetracyclines resistance | tet(L) | | 19,1 (14) | 34,6 (35) | 30,6 (38) | .025 | .07 |
| | | tet(M) | | 46,5 (34) | 44,5 (45) | 60,4 (75) | .385 | .065 |
| | | tet(S) | | 1,3 (1) | 0,9 (1) | 3,2 (4) | .387 | .261 |
| | | Total | | 50,6 (37) | 57,4 (58) | 63,7 (79) | .268 | .078 |
| | Phlopsa resistance | cat | | 9,5 (7) | 0 (0) | 9,6 (12) | .008 | .398 |
| | | fexA | | 0 (0) | 0 (0) | 0,8 (1) | 1 | .241 |
| | | cfrB | | 0 (0) | 0 (0) | 0,8 (1) | 1 | .241 |
| | | optrA | | 0 (0) | 0 (0) | 0,8 (1) | 1 | .241 |
| | Phlopsa resistance | 23s mutation | | 0 (0) | 2,9 (3) | 2,9 (3) | .088 | .062 |
| | | Total | | 9,5 (7) | 2,9 (3) | 11,2 (14) | .089 | .37 |
| Virulence | Pili forming clusters | PilA | SrtA | 79,4 (58) | 87,1 (88) | 85,4 (106) | .164 | .229 |
| | | | Fms21 | 78 (57) | 87,1 (88) | 89,5 (111) | .12 | .047 |
| | | | SrtC4 | 73,9 (54) | 81,1 (82) | 70,9 (88) | .213 | .359 |
| | | | Fms20 | 75,3 (55) | 88,1 (89) | 91,12 (113) | .04 | .007 |
| | | PilB | EbpA | 76,7 (56) | 97 (98) | 97,5 (121) | < .001 | < .001 |
| | | | EbpB | 73,9 (54) | 97 (98) | 97,5 (121) | < .001 | < .001 |
| | | | EbpC | 75,3 (55) | 95 (96) | 98,3 (122) | < .001 | < .001 |
| | | | SrtC1 | 83,5 (61) | 97 (98) | 99,1 (123) | .006 | < .001 |
| | | | Fms14 | 68,4 (50) | 91 (92) | 91,1 (113) | < .001 | < .001 |
| | | | Fms17 | 69,8 (51) | 88,1 (89) | 87 (108) | .005 | .007 |
| | | | Fms13 | 91,7 (67) | 98 (99) | 93,5 (116) | .08 | .358 |
| | | | SrtC2 | 100 (73) | 100 (101) | 100 (124) | 1 | 1 |
| | | | Fms11 | 30,1 (22) | 81,1 (82) | 95,9 (119) | < .001 | < .001 |

| | | | | | | |
|---|---|---|---|---|---|---|
| | Fms19 | 27,3 (20) | 86,1 (87) | 95,9 (119) | < .001 | < .001 |
| | Fms16 | 30,1 (22) | 87,1 (88) | 96,7 (120) | < .001 | < .001 |
| | SrtC3 | 30,1 (22) | 87,1 (88) | 96,7 (120) | < .001 | < .001 |
| LPxTG-containing proteins Adhesins | Acm | 53,4 (39) | 100 (101) | 99,1 (123) | < .001 | < .001 |
| | Scm | 21,9 (16) | 56,4 (57) | 48,38 (60) | < .001 | < .001 |
| | EcbA | 4,1 (3) | 59,4 (60) | 38,7 (48) | < .001 | < .001 |
| | Esp*fm* | 0 (0) | 28,7 (29) | 25 (31) | < .001 | < .001 |
| | SgrA | 6,8 (5) | 80,1 (81) | 66,93 (83) | < .001 | < .001 |
| | Fms3 | 100 (73) | 100 (101) | 98,4 (122) | 1 | .145 |
| | Fms6 | 84,9 (62) | 100 (101) | 99,2 (123) | < .001 | .001 |
| | Fms7 | 83,6 (61) | 96 (97) | 95,2 (118) | .013 | .019 |
| | Fms12 | 93,2 (68) | 98 (99) | 97,6 (121) | .134 | .159 |
| | Fms22 | 24,7 (18) | 0,9 (1) | 58,1 (72) | < .001 | < .001 |
| | Fnm | 100 (100) | 98 (99) | 97,5 (121) | .142 | .081 |
| Secreted antigen A | SagA | 87,6 (64) | 98 (99) | 94,3 (117) | .016 | .124 |
| Carbohydrate transport and gut colonization | PtsD | 5,4 (4) | 95 (96) | 93,5 (116) | < .001 | < .001 |
| Glycosyl hydrolase | *hyl*Efm | 0 (0) | 19,8 (20) | 34,6 (43) | < .001 | < .001 |
| WxL-containing proteins | SwpA | 100 (100) | 100 (101) | 100 (124) | 1 | 1 |
| | SwpB | 94,5 (69) | 97 (98) | 98,3 (122) | .291 | .17 |
| | SwpC | 84,9 (62) | 27,7 (28) | 91,1 (113) | < .001 | .179 |

| | | | | | | | |
|---|---|---|---|---|---|---|---|
| Plasmid families | *pLG1* megaplasmid | rep-Efae287 | 82,1 (60) | 85,1 (86) | 84,6 (105) | .347 | .36 |
| | *pRUM* | rep17 | 10,9 (8) | 79,2 (80) | 62,09 (77) | < .001 | < .001 |
| | Rolling circle replication | rep14 | 6,8 (5) | 18,8 (19) | 46,77 (58) | .019 | < .001 |
| | Rep_3 | rep-pCIZ2 | 6,8 (5) | 20,7 (21) | 37,9 (47) | .008 | < .001 |
| | | rep11 | 4,1 (3) | 28,7 (29) | 20,96 (26) | < .001 | < .001 |
| | | rep18 | 6,8 (5) | 28,7 (29) | 28,22 (35) | < .001 | < .001 |
| | Inc18 (pRE25) | rep2 | 26 (19) | 40,5 (41) | 40,3 (50) | .049 | .042 |
| | | rep1 | 34,2 (25) | 0,9 (1) | 10,48 (13) | < .001 | < .001 |
| | Broad range hosts | rep7 | 1,3 (1) | 0,9 (1) | 4 (5) | .387 | .188 |
| | Described in *S. aureus* | rep22 | 5,4 (4) | 15,8 (16) | 20,16 (25) | .028 | .001 |
| | | rep-pUB110 | 8,2 (6) | 15,8 (16) | 20,16 (25) | .116 | .018 |
| | Rep trans | rep-pHTBeta | 1,3 (1) | 1,9 (2) | 14,51 (18) | .379 | < .001 |
| Insertion Sequences | IS16 | IS16 | 9,5 (7) | 95 (96) | 99,1 (123) | < .001 | < .001 |
| | IS3 | ISefa10, ISEfa8 and ISenfa3 | 75,3 (55) | 96 (97) | 99,19 (123) | < .001 | < .001 |
| | IS1182 | ISefa12 and ISEfa7 | 36,9 (27) | 89,1 (90) | 91,12 (113) | < .001 | < .001 |
| | IS200/IS605 | ISEfa4 and ISEnfa200 | 24,6 (18) | 90 (91) | 74,19 (92) | < .001 | < .001 |
| | IS19/IS982 | ISEfm1 | 91,7 (67) | 80,1 (81) | 90,3 (112) | .03 | .377 |
| | ISL3 | ISEfa11 and ISEfa5 | 20,5 (15) | 82,1 (83) | 82,25 (102) | < .001 | < .001 |
| | IS6 *Lactococcus lactis* | ISS1 | 71,2 (52) | 83,1 (84) | 53,2 (66) | .073 | .013 |
| | IS6 *Lactobacillus sake* | ISS1W | 78 (57) | 29,7 (30) | 37,09 (46) | < .001 | < .001 |
| | IS256 | ISEfa13 and ISEfm2 | 76,7 (56) | 30,6 (31) | 43,5 (54) | < .001 | < .001 |
| | IS256 *Lactococcus garvieae* | ISLgar5 | 26 (19) | 6,9 (7) | 44,3 (55) | 0,001 | .01 |
| | IS5 | ISKpn26 | 0 (0) | 0,9 (1) | 0 (0) | .252 | 1 |
| CRISPR | | | 38,3 (28) | 23,7 (24) | 30,6 (38) | .048 | .219 |
| Cas-system | | | 0 (0) | 0 (0) | 1,6 (2) | 1 | .145 |

| | | | PBP5 amino acid positions | | | | | | | | | | | | | | | | | | | | | | | | | | | | | | | | | | | | | | | | | | | | | | | | | | | | | | | | | | | | | | | | | | | | | | | | | | | | | | | | | | | | | | | | | | | | | | | | | | | | | | | | | | | | | | | |
| --- | --- | --- | --- | --- | --- | --- | --- | --- | --- | --- | --- | --- | --- | --- | --- | --- | --- | --- | --- | --- | --- | --- | --- | --- | --- | --- | --- | --- | --- | --- | --- | --- | --- | --- | --- | --- | --- | --- | --- | --- | --- | --- | --- | --- | --- | --- | --- | --- | --- | --- | --- | --- | --- | --- | --- | --- | --- | --- | --- | --- | --- | --- | --- | --- | --- | --- | --- | --- | --- | --- | --- | --- | --- | --- | --- | --- | --- | --- | --- | --- | --- | --- | --- | --- | --- | --- | --- | --- | --- | --- | --- | --- | --- | --- | --- | --- | --- | --- | --- | --- | --- | --- | --- | --- | --- | --- | --- | --- | --- | --- | --- | --- | --- | --- |
| | | | 2 | 3 | 4 | 5 | 17 | 20 | 24 | 25 | 27 | 28 | 29 | 32 | 34 | 39 | 60 | 66 | 67 | 68 | 73 | 85 | 94 | 95 | 100 | 111 | 115 | 141 | 144 | 151 | 159 | 172 | 177 | 204 | 216 | 218 | 226 | 235 | 262 | 265 | 271 | 272 | 276 | 280 | 290 | 295 | 301 | 314 | 317 | 318 | 324 | 325 | 326 | 339 | 344 | 346 | 349 | 350 | 354 | 357 | 358 | 386 | 397 | 398 | 401 | 405 | 406 | 407 | 408 | 414 | 426 | 430 | 434 | 439 | 440 | 451 | 460 | 461 | 462 | 466 | 466 | 470 | 476 | 485 | 496 | 497 | 499 | 509 | 510 | 524 | 525 | 546 | 552 | 553 | 558 | 561 | 562 | 573 | 581 | 582 | 586 | 598 | 601 | 606 | 625 | 626 | 629 | 632 | 642 | 644 | 646 | 663 | 667 | 672 |
| Previously described by Pietta et al. AAC. 2014. | | | | | | | | | X | | X | | | | X | | | X | | X | | X | | | X | | | | X | | | X | X | X | X | | | | | | | | | | | | | | | | X | | | | | | | | | | | | | | | | | | | | | | | | | | | | | | | | X | X | | | X | X | | X | | | | X | | | | | | | | | | X | | | | | | X | | | | | | X |
| Used in prediction model | | | | | | | | | X | | X | | | | X | X | | X | | X | | X | | | X | | | | X | | | X | X | X | | | | X | | | | | | | | | | | | | X | | | | | | | | | | | | | X | | | | | | | | | X | | | | X | X | X | | | | | X | X | X | | | X | | X | | | | X | | | X | X | X | X | | | | | | X | X | X | | X | | X | |
| | AMP | Accession | | | | | | | | | | | | | | | | | | | | | | | | | | | | | | | | | | | | | | | | | | | | | | | | | | | | | | | | | | | | | | | | | | | | | | | | | | | | | | | | | | | | | | | | | | | | | | | | | | | | | | | | | | | | | | | | | | |
| Com15 | 0,25 | ACBD01000014.1 | K | R | S | D | A | V | V | A | S | G | G | Y | R | T | D | G | M | A | A | E | A | D | E | D | S | N | K | P | E | T | L | D | A | D | Q | P | G | Y | A | A | G | D | P | N | S | T | G | K | T | D | G | N | K | T | A | K | T | D | S | D | E | D | A | N | P | E | Q | F | M | A | D | D | P | W | Y | Q | V | - | S | Q | A | M | N | F | A | D | L | E | E | N | A | M | A | G | T | E | I | G | V | Q | N | S | E | K | E | K | P | N | E | T | P | Y |
| E0120 | 0,5 | AHWI01000003.1 | . | . | . | . | . | . | A | . | G | . | . | . | . | . | . | . | . | . | . | . | . | . | Q | . | . | . | Q | . | . | A | . | . | . | . | . | T | . | . | . | . | . | . | . | . | . | . | . | . | . | . | . | . | . | . | . | . | . | . | . | . | . | . | . | . | . | . | . | . | . | . | . | N | . | . | . | . | . | . | . | . | . | . | K | . | T | . | F | . | D | . | . | . | . | . | . | D | V | S | . | . | . | . | . | . | . | . | . | . | G | . | . | . |
| EnGen0048 | 0,5 | AHYE01000001.1 | . | . | . | . | . | . | A | . | G | . | . | . | Q | . | . | E | . | . | . | . | . | . | Q | . | . | . | Q | . | . | A | I | . | S | . | . | . | . | . | . | . | . | . | . | . | . | . | . | . | A | . | . | . | . | . | . | . | . | . | N | . | . | . | S | . | . | . | . | . | . | . | . | . | . | . | . | . | . | . | . | . | . | . | K | . | I | . | . | . | D | . | . | . | . | . | . | . | . | . | . | . | . | . | . | . | . | . | . | . | G | . | . | . |
| UC7267 | 0,5 | ASAM01000001.1 | . | . | . | . | . | . | A | . | G | . | . | . | Q | . | . | E | . | . | . | . | . | . | Q | . | . | . | Q | . | . | A | I | . | S | . | . | . | . | . | . | . | . | . | . | . | . | . | . | . | A | . | . | . | . | . | . | . | . | . | . | . | . | . | S | . | . | . | . | . | . | . | . | . | . | . | . | . | . | . | . | . | . | . | K | . | I | . | . | . | D | . | . | . | . | . | . | . | . | . | . | . | . | . | . | . | . | . | . | . | G | . | . | . |
| E980 | 1 | ABQA01000020.1 | . | . | . | . | . | . | A | . | G | . | . | . | . | . | . | . | . | . | T | . | . | . | . | . | . | . | Q | . | . | . | . | . | . | . | . | . | . | . | . | . | . | . | . | . | . | . | . | . | A | . | . | . | . | . | . | . | . | . | . | . | . | . | . | . | . | . | . | . | . | . | . | . | . | . | . | . | . | . | . | . | . | . | . | . | . | . | . | . | . | . | . | . | . | . | . | . | . | . | . | . | . | . | . | . | . | . | . | . | G | . | . | . |
| TX1330 | 1 | ACHL01000102.1, ACHL01000101.1, EEI60277 | . | . | . | . | . | . | . | . | . | . | . | . | . | . | . | . | . | . | . | . | . | . | . | . | . | . | . | . | . | . | . | . | . | . | . | . | . | . | . | . | . | . | . | . | . | . | . | . | . | . | . | . | . | . | . | . | . | . | . | . | . | . | . | X | X | . | . | . | . | . | . | . | . | . | . | . | . | . | . | . | . | . | . | . | . | . | . | . | . | . | . | . | . | . | . | . | . | . | . | . | . | . | . | . | . | . | . | . | G | . | A | . |
| E1185 | 1 | AHWS01000017.1 | . | . | . | . | . | . | A | . | G | . | . | . | Q | . | . | E | . | . | . | . | . | . | Q | . | . | . | Q | . | . | A | I | . | S | . | . | . | . | . | . | . | . | . | . | . | . | . | . | . | A | . | . | . | . | . | . | . | . | . | . | . | . | . | S | . | . | . | . | . | . | . | . | . | . | . | . | . | . | . | . | . | . | . | K | . | I | . | . | . | D | . | . | . | . | . | . | . | . | . | . | . | . | . | . | . | . | . | . | . | G | . | . | . |
| E0164 | 1 | AHWJ01000010.1 | . | . | . | . | S | . | A | . | G | . | . | . | Q | N | . | E | . | . | . | . | . | . | Q | . | . | . | Q | . | . | A | I | . | S | . | . | . | . | . | . | . | . | . | . | . | . | . | . | . | A | . | . | . | . | . | . | . | . | . | . | . | . | . | . | . | . | . | . | . | . | . | . | . | . | . | . | . | . | . | . | . | . | . | K | . | T | . | . | . | D | . | . | . | . | . | . | . | . | . | . | . | . | . | . | . | . | . | . | . | G | . | . | . |
| E1627 | 1 | AHXJ01000003.1 | . | . | . | . | . | . | A | . | G | . | . | . | Q | N | . | E | . | . | . | . | . | . | Q | . | . | . | Q | . | . | A | I | . | S | . | . | . | . | . | . | . | . | . | . | . | . | . | . | . | A | . | . | . | . | . | . | . | . | . | . | . | . | . | . | . | . | . | . | . | . | . | . | . | . | . | . | . | . | . | . | . | . | . | K | . | T | . | . | . | D | . | . | . | . | . | . | . | . | . | . | . | . | . | . | . | . | . | . | . | G | . | . | . |
| UC10237 | 1 | AWWP01000001.1 | . | . | . | . | . | . | A | . | G | . | . | . | Q | . | . | E | . | T | . | D | . | . | Q | . | . | . | Q | . | . | A | I | G | S | . | . | . | . | . | . | . | . | . | . | . | . | . | . | . | A | . | . | . | . | . | . | . | . | . | . | . | . | . | . | . | . | . | . | . | . | . | . | . | . | . | . | . | . | . | . | . | . | . | K | . | T | . | . | . | D | . | . | . | I | . | C | . | . | . | . | . | . | . | . | . | . | . | . | . | . | . | G | . | S | . |
| E1573 | 1 | AHWX01000001.1 | . | . | . | . | . | . | A | . | G | . | . | . | Q | . | . | E | . | T | . | D | . | . | Q | . | . | . | Q | . | . | A | I | G | S | . | . | . | . | . | . | . | . | . | . | . | . | . | . | . | A | . | . | . | . | . | . | . | . | . | . | . | . | . | . | . | . | . | . | . | . | . | . | . | . | . | . | . | . | . | . | . | . | . | K | . | T | . | . | . | D | . | . | . | . | . | . | . | . | . | . | . | . | . | . | . | . | . | . | . | G | . | S | . |
| UC7256 | 2 | AWWM01000001.1 | . | . | . | . | . | . | A | . | G | . | . | . | . | . | . | . | . | . | T | . | . | . | . | . | . | . | Q | . | . | . | . | . | . | . | . | . | . | . | . | . | . | . | . | . | . | . | . | . | A | . | . | . | . | . | . | . | . | E | . | . | G | . | . | . | . | . | . | . | . | . | . | . | . | . | . | . | . | . | . | . | . | . | . | . | . | . | . | . | . | . | . | . | . | . | . | . | . | . | . | . | . | . | . | . | . | . | . | . | G | . | . | . |
| 1141733 | 2 | ACAZ01000019.1 | . | . | . | . | . | . | . | . | . | . | . | . | . | . | . | . | . | . | . | . | . | . | . | . | . | . | . | . | . | . | . | . | . | . | . | . | . | . | . | . | . | . | L | . | . | . | . | . | . | . | . | . | . | . | . | . | . | . | . | . | . | . | . | . | . | . | . | . | . | . | . | . | . | . | . | . | . | . | . | . | . | . | . | . | . | . | . | . | . | . | . | . | . | . | . | . | . | . | . | . | . | . | . | . | . | . | . | . | G | . | L | . |
| UC8733 | 2 | AWWO01000001.1 | . | . | . | . | . | . | A | . | G | . | . | . | Q | N | . | E | . | . | . | . | . | . | Q | . | . | . | Q | . | . | A | I | . | S | . | . | . | . | . | . | . | . | . | . | . | . | I | . | . | A | . | . | . | . | . | . | . | . | . | . | . | . | . | . | . | A | . | . | . | . | . | . | . | . | . | . | . | . | . | . | . | . | . | K | . | T | . | . | . | D | . | . | . | . | . | . | . | . | . | . | . | . | . | . | . | . | . | . | . | G | . | . | . |
| E0680 | 4 | AHWN01000006.1 | . | . | . | Y | . | . | A | . | G | . | . | . | Q | . | . | E | . | . | . | . | . | . | Q | . | . | . | Q | . | . | A | I | . | S | . | . | . | . | . | . | . | . | . | . | . | . | . | . | . | A | . | . | . | . | . | . | . | . | . | . | . | . | . | S | . | . | . | . | . | . | . | . | . | . | . | . | . | . | . | . | . | . | . | K | . | I | . | . | . | D | . | . | . | . | . | . | . | . | . | . | . | . | . | . | . | . | . | . | . | G | . | . | . |
| E0045 | 4 | AHWH01000023.1 | . | . | . | . | . | . | A | . | G | . | . | . | Q | . | . | E | . | . | . | . | . | . | Q | . | . | . | Q | . | . | A | I | . | S | . | . | . | . | . | . | . | . | . | . | . | . | . | . | . | A | . | . | . | . | . | . | . | . | . | . | . | . | . | S | . | . | . | . | . | . | . | . | . | . | . | . | . | . | . | . | . | . | . | K | . | I | . | . | . | D | . | . | . | . | . | . | . | . | . | . | . | . | . | . | . | . | . | . | . | G | . | . | . |
| E1552 | 8 | AHWW01000007.1 | . | . | . | . | . | . | A | . | G | . | . | . | Q | . | . | E | . | . | . | . | . | . | Q | . | . | . | Q | . | . | A | I | . | S | . | . | . | . | . | . | . | . | . | . | . | . | . | . | . | A | . | . | . | . | . | . | . | . | . | . | . | . | . | S | . | . | . | . | . | . | . | . | . | . | . | . | . | . | . | . | . | . | . | K | . | I | . | . | . | D | . | . | . | . | . | . | . | . | . | . | . | . | . | . | . | . | . | . | . | G | . | . | . |
| E1613 | ≤8 | AHXE01000011.1 | . | . | . | . | . | . | . | . | G | . | S | . | . | . | . | . | . | . | . | . | . | N | . | . | . | . | . | . | . | . | . | . | . | G | . | H | . | . | . | . | . | . | . | S | . | . | . | . | A | . | . | . | . | . | . | . | . | . | . | . | . | . | . | . | . | . | . | . | . | . | . | . | . | . | . | . | . | . | . | . | . | . | . | . | . | . | . | . | . | . | . | . | . | . | . | . | . | . | . | . | . | . | . | . | . | . | . | . | . | G | . | . | . |
| E1861 | ≤8 | AHXP01000002.1 | . | . | . | . | . | . | A | . | G | . | . | . | . | . | . | . | . | . | T | . | . | . | . | . | . | . | Q | . | . | . | . | . | . | . | . | . | . | . | . | . | . | . | . | . | . | . | . | R | A | . | . | . | . | . | . | . | . | . | . | . | . | . | . | . | . | . | . | . | . | . | . | . | . | . | . | . | . | . | . | . | . | . | . | . | . | . | . | . | . | . | . | . | . | . | . | . | . | . | . | . | . | . | . | . | . | . | . | . | G | . | . | . |
| E2620 | ≤8 | AHXW01000008.1 | . | . | . | . | . | . | . | . | G | . | . | . | . | . | . | . | . | . | . | . | . | . | . | . | . | . | . | L | . | . | . | . | V | . | . | . | . | . | . | . | . | . | . | . | . | . | . | . | . | . | . | . | . | . | . | . | . | . | . | . | . | . | . | . | . | . | . | . | . | . | . | . | . | . | . | . | . | . | . | . | . | . | . | . | . | . | . | G | . | . | . | . | . | . | . | . | . | . | . | . | . | . | . | . | . | . | . | . | . | . | G | . | . | . |
| BM4107 | ≤8 | AF364092.1 | . | . | . | . | . | . | . | T | . | . | . | . | . | S | . | . | . | . | . | . | . | . | . | . | . | . | . | . | . | . | . | . | . | . | . | . | . | . | . | . | . | . | . | . | . | . | . | . | . | . | . | . | . | . | . | . | . | . | . | . | . | . | . | . | . | . | . | . | . | . | . | . | . | . | . | . | . | . | . | . | . | . | . | . | . | . | . | . | . | . | . | . | . | . | . | . | . | . | . | . | . | . | . | . | . | . | . | D | G | . | . | . |
| E1604 | ≤8 | AHXD01000013.1 | . | . | . | . | . | . | . | . | . | . | . | . | . | . | . | . | . | . | . | . | . | . | . | . | . | . | . | . | . | . | . | . | . | . | . | . | . | . | . | . | . | . | . | . | . | . | . | . | . | . | . | . | . | . | . | . | . | . | . | N | . | . | . | . | . | . | . | . | . | . | . | . | . | - | . | . | . | . | . | . | . | . | . | . | . | . | . | . | . | . | . | . | . | . | . | . | . | . | . | . | . | . | . | . | . | . | . | . | G | . | . | . |
| E3548 | ≤8 | AHYB01000002.1 | . | . | . | . | . | A | . | . | . | . | . | . | . | . | . | . | . | . | . | . | . | . | . | . | . | . | . | . | . | . | . | . | . | . | . | . | . | . | . | . | . | . | . | . | . | . | . | . | . | . | . | . | . | . | . | . | . | . | . | . | . | . | . | . | . | . | . | . | . | . | . | . | . | . | . | . | . | . | . | . | . | . | . | . | . | . | . | . | . | . | . | . | . | . | . | . | . | . | . | . | . | . | . | . | . | . | . | . | G | . | . | . |
| D63r | ≤8 | X84860.1 | . | . | . | . | . | . | . | . | . | . | . | . | . | . | . | . | . | . | . | . | . | . | . | . | . | . | . | . | . | . | . | . | . | . | . | . | . | . | . | . | . | . | . | . | . | . | . | . | . | . | . | . | . | . | . | . | . | . | . | . | . | . | . | . | . | . | . | . | . | . | . | . | . | . | . | . | . | . | . | H | . | . | . | . | . | . | . | . | . | . | . | . | . | . | . | . | . | . | . | . | . | . | . | . | . | . | . | . | G | . | . | . |
| 1231501 | ≤8 | ACAY01000033.1 | . | . | . | . | . | . | . | . | . | . | . | . | . | . | . | . | . | . | . | . | . | . | . | . | . | . | . | . | . | . | . | . | . | . | . | . | . | . | . | . | . | . | . | . | . | . | . | . | . | . | . | . | . | . | . | . | . | . | . | . | . | . | . | . | . | . | . | . | . | . | . | . | . | . | . | . | . | . | . | . | . | . | . | . | . | . | . | . | . | . | . | . | . | . | . | . | . | . | . | . | . | . | . | . | . | . | . | . | G | . | . | . |
| E1007 | ≤8 | AHWP01000007.1 | . | . | . | . | . | . | . | . | . | . | . | . | . | . | . | . | . | . | . | . | . | . | . | . | . | . | . | . | . | . | . | . | . | . | . | . | . | . | . | . | . | . | . | . | . | . | . | . | . | . | . | . | . | . | . | . | . | . | . | . | . | . | . | . | . | . | . | . | . | . | . | . | . | . | . | . | . | . | . | . | . | . | . | . | . | . | . | . | . | . | . | . | . | . | . | . | . | . | . | . | . | . | . | . | . | . | . | . | G | . | . | . |
| E3083 | ≤8 | AHXZ01000002.1 | . | . | . | . | . | . | . | . | . | . | . | . | . | . | . | . | . | . | . | . | . | . | . | . | . | . | . | . | . | . | . | . | . | . | . | . | . | . | . | . | . | . | . | . | . | . | . | . | . | . | . | . | . | . | . | . | . | . | . | . | . | . | . | . | . | . | . | . | . | . | . | . | . | . | . | . | . | . | . | . | . | . | . | . | . | . | . | . | . | . | . | . | . | . | . | . | . | . | . | . | . | . | . | . | . | . | . | . | G | . | . | . |
| E1590 | ≤8 | AHXC01000004.1 | . | . | . | . | . | . | . | . | . | . | . | . | . | . | . | . | . | . | . | . | . | . | . | . | . | . | . | . | . | . | . | . | . | . | . | . | . | . | . | . | . | . | . | . | . | . | . | . | . | . | . | . | . | . | . | . | . | . | . | . | . | . | . | . | . | . | . | . | . | . | . | . | . | . | . | . | . | . | . | . | . | . | . | . | . | . | . | . | . | . | . | . | . | . | . | . | . | . | . | . | . | . | . | . | . | . | . | . | G | . | L | . |
| Com12 | ≤8 | ACBC01000012.1 | . | . | . | . | . | . | . | . | . | . | . | . | . | . | . | . | . | . | . | . | . | . | . | . | . | . | . | . | . | . | . | . | . | . | . | . | . | . | . | . | . | . | . | . | . | . | . | . | . | . | . | . | . | . | . | . | . | . | . | . | . | . | . | . | . | . | . | . | . | . | . | . | . | . | . | . | . | . | . | . | . | . | . | . | . | . | . | . | . | . | . | . | . | . | . | . | . | . | . | . | . | . | . | . | . | . | . | . | G | . | A | . |
| VAN327 | ≤8 | ASEC01000004.1 | . | . | . | . | . | . | A | . | G | . | . | . | . | . | . | . | . | . | . | . | . | . | Q | . | . | . | Q | . | . | A | . | . | . | . | . | T | . | . | . | . | . | . | . | . | . | . | . | . | . | . | S | . | . | . | . | . | . | . | . | . | . | . | . | . | . | . | . | . | . | . | . | N | . | . | . | . | . | . | . | . | . | . | K | . | T | . | F | . | D | . | . | . | . | . | . | D | V | S | . | . | . | . | . | . | . | . | . | . | G | . | . | . |
| VAN335 | ≤8 | AIUZ01000021.1 | . | . | . | . | . | . | A | . | G | . | . | . | . | . | . | . | . | . | . | . | . | . | Q | . | . | . | Q | . | . | A | . | . | . | . | . | T | . | . | . | . | . | . | . | . | . | . | . | . | . | . | S | . | . | . | . | . | . | . | . | . | . | . | . | . | . | . | . | . | . | . | . | N | . | . | . | . | . | . | . | . | . | . | K | . | T | . | F | . | D | . | . | . | . | . | . | D | V | S | . | . | . | . | . | . | . | . | . | . | G | . | . | . |
| E1574 | ≤8 | AHWY01000003.1 | . | . | . | . | . | . | A | . | G | . | . | . | . | . | . | . | . | . | . | . | . | . | Q | . | . | . | Q | . | . | A | . | . | . | . | . | T | . | . | . | . | . | . | . | . | . | . | . | . | . | . | . | . | . | . | . | . | . | . | . | . | . | . | . | . | . | . | . | . | . | . | . | N | . | . | . | . | . | . | . | . | . | . | K | . | T | . | F | . | D | . | . | . | . | . | . | D | V | S | . | . | . | . | . | . | . | . | . | . | G | . | . | . |
| E2071 | ≤8 | AHXT01000013.1 | . | . | . | . | . | . | A | . | G | . | . | . | . | . | . | . | . | . | . | . | . | . | Q | . | . | . | Q | . | . | A | . | . | . | . | . | T | . | . | . | . | . | . | . | . | . | . | . | . | . | . | . | . | . | . | . | . | . | . | . | . | . | . | . | . | . | . | . | . | . | . | . | N | . | . | . | . | . | . | . | . | . | . | K | . | T | . | F | . | D | . | . | . | . | . | . | D | V | S | . | . | . | . | . | . | . | . | . | . | G | . | . | . |
| D366 | ≤8 | X84859.1 | - | - | - | - | . | . | A | . | G | . | . | . | Q | . | . | E | . | . | . | . | . | . | Q | . | . | . | Q | . | . | A | I | G | S | . | . | . | . | . | . | . | . | . | . | . | . | . | . | . | A | . | . | . | . | . | . | . | . | . | . | . | . | . | S | . | . | . | . | . | . | . | . | . | . | . | . | . | . | . | . | . | . | . | K | . | I | . | . | . | D | . | . | . | . | . | . | . | . | . | . | R | . | . | . | . | . | . | . | . | G | . | . | . |
| E1039 | ≤8 | ACOS01000009.1 | . | . | . | . | . | . | A | . | G | . | . | . | Q | . | . | E | . | . | . | . | . | . | Q | . | . | . | Q | . | . | A | I | | S | . | . | . | . | . | . | . | . | . | . | . | . | . | . | . | A | . | . | . | . | . | . | . | . | . | . | . | . | N | S | . | . | . | . | . | . | . | . | . | . | . | . | . | . | . | . | . | . | . | K | . | I | . | . | . | D | . | . | . | . | . | . | . | . | . | . | . | . | . | . | . | . | . | . | . | G | . | . | . |
| E1634 | ≤8 | AHXL01000003.1 | . | . | . | . | . | . | A | . | G | . | . | . | Q | . | . | E | . | . | . | . | . | . | Q | . | . | . | Q | . | . | A | I | | S | . | . | . | . | . | . | . | . | . | . | . | . | . | . | . | A | . | . | . | . | . | . | . | . | . | . | . | . | . | S | . | . | K | . | . | . | . | . | . | . | . | . | . | . | . | . | . | . | . | K | . | I | . | . | . | D | . | . | . | . | . | . | . | . | . | . | . | . | . | . | . | . | . | . | . | G | . | . | . |
| 841V03 | ≤8 | AISU01000016.1 | . | . | . | . | . | . | A | . | G | . | . | . | Q | . | . | E | . | . | . | . | . | . | Q | . | . | . | Q | . | . | A | I | | S | . | . | . | . | . | . | . | . | . | . | . | . | . | . | . | A | . | . | . | . | . | . | . | . | . | . | . | . | . | S | . | . | . | . | . | . | . | . | . | . | . | . | . | . | . | . | . | . | . | K | . | I | . | . | . | D | . | . | . | . | . | . | . | . | . | . | . | . | . | . | . | . | . | . | . | G | . | . | . |
| E0679 | ≤8 | AHWM01000004.1 | . | . | . | . | . | . | A | . | G | . | . | . | Q | . | . | E | . | . | . | . | . | . | Q | . | . | . | Q | . | . | A | I | | S | . | . | . | . | . | . | . | . | . | . | . | . | . | . | . | A | . | . | . | . | . | . | . | . | . | . | . | . | . | S | . | . | . | . | . | . | . | . | . | . | . | . | . | . | . | . | . | . | . | K | . | I | . | . | . | D | . | . | . | . | . | . | . | . | . | . | . | . | . | . | . | . | . | . | . | G | . | . | . |
| E1575 | ≤8 | AHWZ01000016.1 | . | . | . | . | . | . | A | . | G | . | . | . | Q | . | . | E | . | . | . | . | . | . | Q | . | . | . | Q | . | . | A | I | | S | . | . | . | . | . | . | . | . | . | . | . | . | . | . | . | A | . | . | . | . | . | . | . | . | . | . | . | . | . | S | . | . | . | . | . | . | . | . | . | . | . | . | . | . | . | . | . | . | . | K | . | I | . | . | . | D | . | . | . | . | . | . | . | . | . | . | . | . | . | . | . | . | . | . | . | G | . | . | . |
| E1620 | ≤8 | AHXF01000060.1 | . | . | . | . | . | . | A | . | G | . | . | . | Q | . | . | E | . | . | . | . | . | . | Q | . | . | . | Q | . | . | A | I | | S | . | . | . | . | . | . | . | . | . | . | . | . | . | . | . | A | . | . | . | . | . | . | . | . | . | . | . | . | . | S | . | . | . | . | . | . | . | . | . | . | . | . | . | . | . | . | . | . | . | K | . | I | . | . | . | D | . | . | . | . | . | . | . | . | . | . | . | . | . | . | . | . | . | . | . | G | . | . | . |
| HF50104 | ≤8 | AITR01000020.1 | . | . | . | . | . | . | A | . | G | . | . | . | Q | . | . | E | . | . | . | . | . | . | Q | . | . | . | Q | . | . | A | I | | S | . | . | . | . | . | . | . | . | . | . | . | . | . | . | . | A | . | . | . | . | . | . | . | . | . | . | . | . | . | S | . | . | . | . | . | . | . | . | . | . | . | . | . | . | . | . | . | . | . | K | . | I | . | . | . | D | . | . | . | . | . | . | . | . | . | . | . | . | . | . | . | . | . | . | . | G | . | . | . |
| HF50105 | ≤8 | AITS01000013.1 | . | . | . | . | . | . | A | . | G | . | . | . | Q | . | . | E | . | . | . | . | . | . | Q | . | . | . | Q | . | . | A | I | | S | . | . | . | . | . | . | . | . | . | . | . | . | . | . | . | A | . | . | . | . | . | . | . | . | . | . | . | . | . | S | . | . | . | . | . | . | . | . | . | . | . | . | . | . | . | . | . | . | . | K | . | I | . | . | . | D | . | . | . | . | . | . | . | . | . | . | . | . | . | . | . | . | . | . | . | G | . | . | . |
| HF50106 | ≤8 | AITT01000026.1 | . | . | . | . | . | . | A | . | G | . | . | . | Q | . | . | E | . | . | . | . | . | . | Q | . | . | . | Q | . | . | A | I | | S | . | . | . | . | . | . | . | . | . | . | . | . | . | . | . | A | . | . | . | . | . | . | . | . | . | . | . | . | . | S | . | . | . | . | . | . | . | . | . | . | . | . | . | . | . | . | . | . | . | K | . | I | . | . | . | D | . | . | . | . | . | . | . | . | . | . | . | . | . | . | . | . | . | . | . | G | . | . | . |
| HF50203 | ≤8 | AITU01000017.1 | . | . | . | . | . | . | A | . | G | . | . | . | Q | . | . | E | . | . | . | . | . | . | Q | . | . | . | Q | . | . | A | I | | S | . | . | . | . | . | . | . | . | . | . | . | . | . | . | . | A | . | . | . | . | . | . | . | . | . | . | . | . | . | S | . | . | . | . | . | . | . | . | . | . | . | . | . | . | . | . | . | . | . | K | . | I | . | . | . | D | . | . | . | . | . | . | . | . | . | . | . | . | . | . | . | . | . | . | . | G | . | . | . |
| HF50215 | ≤8 | AITW01000030.1 | . | . | . | . | . | . | A | . | G | . | . | . | Q | . | . | E | . | . | . | . | . | . | Q | . | . | . | Q | . | . | A | I | | S | . | . | . | . | . | . | . | . | . | . | . | . | . | . | . | A | . | . | . | . | . | . | . | . | . | . | . | . | . | S | . | . | . | . | . | . | . | . | . | . | . | . | . | . | . | . | . | . | . | K | . | I | . | . | . | D | . | . | . | . | . | . | . | . | . | . | . | . | . | . | . | . | . | . | . | G | . | . | . |
| VAN219 | ≤8 | AIUW01000023.1 | . | . | . | . | . | . | A | . | G | . | . | . | Q | . | . | E | . | . | . | . | . | . | Q | . | . | . | Q | . | . | A | I | G | S | . | . | . | . | . | . | . | . | . | . | . | . | . | . | . | A | . | . | . | . | . | . | . | . | . | . | . | . | . | . | . | . | . | . | . | . | . | . | . | . | . | . | . | . | . | . | . | . | . | K | . | T | . | . | . | D | . | . | . | . | . | . | . | . | . | . | . | . | . | . | . | . | . | . | . | G | . | . | . |
| VAN222 | ≤8 | AIUX01000013.1 | . | . | . | . | . | . | A | . | G | . | . | . | Q | . | . | E | . | . | . | . | . | . | Q | . | . | . | Q | . | . | A | I | G | S | . | . | . | . | . | . | . | . | . | . | . | . | . | . | . | A | . | . | . | . | . | . | . | . | . | . | . | . | . | . | . | . | . | . | . | . | . | . | . | . | . | . | . | . | . | . | . | . | . | K | . | T | . | . | . | D | . | . | . | . | . | . | . | . | . | . | . | . | . | . | . | . | . | . | . | G | . | . | . |
| VAN476 | ≤8 | ASEB01000017.1 | . | . | . | . | . | . | A | . | G | . | . | . | Q | . | . | E | . | . | . | . | . | . | Q | . | . | . | Q | . | . | A | I | G | S | . | . | . | . | . | . | . | . | . | . | . | . | . | . | . | A | . | . | . | . | . | . | . | . | . | . | . | . | . | . | . | . | . | . | . | . | . | . | . | . | . | . | . | . | . | . | . | . | . | K | . | T | . | . | . | D | . | . | . | . | . | . | . | . | . | . | . | . | . | . | . | . | . | . | . | G | . | . | . |
| E1258 | ≤8 | AHWT01000005.1 | . | . | . | . | . | . | A | . | G | S | . | . | Q | N | . | E | . | . | . | . | . | . | Q | . | . | . | Q | . | . | A | I | | S | . | . | . | . | . | . | . | . | . | . | . | . | . | . | . | A | . | . | . | . | . | . | . | . | . | . | . | . | . | . | . | . | . | . | . | . | . | E | . | . | . | . | . | . | . | . | . | . | . | K | . | T | . | . | . | D | . | . | . | . | . | . | . | . | . | . | . | . | . | . | . | . | . | . | . | G | . | . | . |
| E0269 | ≤8 | AHWK01000016.1, AHWK0100001.1 | . | . | . | . | S | . | A | . | G | . | . | . | Q | N | . | E | . | . | . | . | . | . | Q | . | . | . | Q | . | . | A | I | | S | . | . | . | . | . | . | . | . | . | . | . | . | . | . | . | A | . | . | . | . | . | . | . | . | . | . | . | . | . | . | . | . | . | . | . | . | . | . | . | . | . | . | . | . | . | . | . | . | . | K | . | T | . | . | . | D | . | . | . | . | . | . | . | . | . | . | . | . | . | . | . | . | . | . | . | G | . | . | . |
| E1630 | ≤8 | AHXK01000019.1 | . | . | . | . | . | . | A | . | G | . | . | . | Q | N | . | E | . | . | . | . | . | . | Q | . | . | . | Q | . | . | A | I | | S | . | . | . | . | . | . | . | . | . | . | . | . | I | . | . | A | . | . | . | . | . | . | . | . | . | . | . | . | . | . | . | A | . | . | . | . | . | . | . | . | . | . | . | . | . | . | . | . | . | K | . | T | E | . | . | D | . | . | . | . | . | . | . | . | . | . | . | . | F | . | . | . | . | . | . | G | . | . | . |
| E1622 | ≤8 | AHXG01000003.1 | . | . | . | . | . | . | A | . | G | . | . | . | Q | . | . | E | . | T | . | D | . | . | Q | . | . | . | Q | . | . | A | I | G | S | . | . | . | . | . | . | . | . | . | . | . | . | . | . | . | A | . | . | . | . | . | . | . | . | . | . | . | . | . | . | . | . | . | . | . | . | . | . | . | . | . | . | . | . | . | . | . | . | . | K | . | T | . | . | . | D | . | . | I | . | . | . | . | . | . | . | . | . | . | . | . | . | . | . | . | G | . | S | . |
| E1071 | ≤8 | ABQI01000087.1 | . | . | . | . | . | . | A | T | G | . | . | . | Q | . | . | E | . | T | . | D | . | . | Q | . | . | . | Q | . | . | A | I | G | S | . | . | . | . | . | . | . | . | . | . | . | . | . | . | . | A | . | . | . | . | . | . | . | . | . | . | . | . | . | . | . | . | . | . | . | . | . | . | . | . | . | . | . | . | . | . | . | . | . | K | . | T | . | . | . | D | . | . | . | . | . | . | . | . | . | . | . | . | . | . | . | . | . | . | . | G | . | S | . |
| E1050 | ≤8 | AHWQ01000005.1 | . | . | . | . | . | . | A | . | G | . | . | . | Q | . | . | E | I | T | . | D | . | . | Q | . | . | . | Q | . | . | A | I | G | S | . | . | . | . | . | . | . | . | . | . | . | . | . | . | . | A | . | . | . | . | . | . | . | . | . | . | . | . | . | . | . | . | . | . | . | . | . | . | . | . | . | . | . | . | . | . | . | . | . | K | . | T | . | . | . | D | . | . | . | . | . | . | . | . | . | . | . | . | . | . | . | . | . | . | . | G | . | S | . |
| E1576 | ≤8 | AHXA01000008.1 | . | . | . | . | . | . | A | . | G | . | . | . | Q | . | . | E | . | T | . | D | . | . | Q | . | . | . | Q | . | . | A | I | G | S | . | . | . | . | . | . | . | . | . | . | . | . | . | . | . | A | . | . | . | . | . | . | . | . | . | . | . | . | . | . | . | . | . | . | . | . | . | . | . | . | . | . | . | . | . | . | . | . | . | K | . | T | . | . | . | D | . | . | . | . | . | . | . | . | . | . | . | . | . | . | . | . | . | . | . | G | . | S | . |
| E1578 | ≤8 | AHXB01000009.1 | . | . | . | . | . | . | A | . | G | . | . | . | Q | . | . | E | . | T | . | D | . | . | Q | . | . | . | Q | . | . | A | I | G | S | . | . | . | . | . | . | . | . | . | . | . | . | . | . | . | A | . | . | . | . | . | . | . | . | . | . | . | . | . | . | . | . | . | . | . | . | . | . | . | . | . | . | . | . | . | . | . | . | . | K | . | T | . | . | . | D | . | . | . | . | . | . | . | . | . | . | . | . | . | . | . | . | . | . | . | G | . | S | . |
| E1623 | ≤8 | AHXH01000005.1 | . | . | . | . | . | . | A | . | G | . | . | . | Q | . | . | E | . | T | . | D | . | . | Q | . | . | . | Q | . | . | A | I | G | S | . | . | . | . | . | . | . | . | . | . | . | . | . | . | . | A | . | . | . | . | . | . | . | . | . | . | . | . | . | . | . | . | . | . | . | . | . | . | . | . | . | . | . | . | . | . | . | . | . | K | . | T | . | . | . | D | . | . | . | . | . | . | . | . | . | . | . | . | . | . | . | . | . | . | . | G | . | S | . |
| VAN342 | ≤8 | AIVA01000020.1 | . | . | . | . | . | . | A | . | G | . | . | . | Q | . | . | E | . | T | . | D | . | . | Q | . | . | . | Q | . | . | A | I | G | S | . | . | . | . | . | . | . | . | . | . | . | . | . | . | . | A | . | . | . | . | . | . | . | . | . | . | . | . | . | . | . | . | . | . | . | . | . | . | . | . | . | . | . | . | . | . | . | . | . | K | . | T | . | . | . | D | . | . | . | . | . | . | . | . | . | . | . | . | . | . | . | . | . | . | . | G | . | S | . |
| VAN332 | ≤8 | AIUY01000015.1 | . | . | . | . | . | . | A | . | G | . | . | . | Q | . | . | E | . | T | . | D | . | . | Q | . | . | . | Q | . | . | A | I | G | S | . | . | . | . | . | . | . | . | . | . | . | . | . | . | . | A | . | . | . | . | . | . | . | . | . | . | . | . | . | . | . | . | . | . | . | . | . | . | . | . | . | . | . | . | . | . | . | . | . | K | . | T | . | . | . | D | . | . | . | . | . | . | . | . | . | . | . | . | . | . | K | . | . | . | . | . | G | . | S | . |
| VAN345 | ≤8 | AIVB01000020.1 | . | . | . | . | . | . | A | . | G | . | . | . | Q | . | . | E | . | T | . | D | . | . | Q | . | . | . | Q | . | . | A | I | G | S | . | . | . | . | . | . | . | . | . | . | . | . | . | . | . | A | . | . | . | . | . | . | . | . | . | . | . | . | . | . | . | . | . | . | . | . | . | . | . | . | . | . | . | . | . | . | . | . | . | K | . | T | . | . | . | D | . | . | . | . | . | . | . | . | . | . | . | . | . | . | K | . | . | . | . | . | G | . | S | . |
| E0688 | ≤8 | AHWO01000001.1 | . | . | . | . | . | . | A | . | G | . | . | . | Q | . | . | E | . | . | . | . | . | . | Q | . | . | . | Q | . | . | A | I | | S | . | . | . | . | . | . | . | . | . | . | . | . | . | . | . | A | . | . | . | . | . | . | . | . | . | . | . | . | . | S | . | . | . | . | . | . | . | . | . | . | . | . | . | . | . | . | . | . | . | K | . | I | . | . | . | D | . | . | . | . | . | . | . | . | . | . | . | . | . | . | . | . | . | . | . | G | . | . | . |
| NRRLB_2354 | ≤8 | CP004063.1 | . | . | . | . | . | . | A | . | G | . | . | . | Q | . | . | E | . | T | . | D | . | . | Q | . | . | . | Q | . | . | A | I | G | S | . | . | . | . | . | . | . | S | . | . | . | . | . | . | . | A | . | . | . | . | . | . | . | . | . | . | . | . | . | . | . | . | . | . | . | . | . | . | . | . | . | . | . | . | . | . | . | . | . | K | . | T | . | . | . | D | . | . | . | . | . | . | . | . | . | . | . | . | . | . | . | . | . | . | . | G | . | S | . |
| E2369 | ≥16 | AHYH01000012.1 | . | . | . | . | . | . | A | . | G | . | . | . | Q | . | . | E | . | T | . | D | . | . | Q | . | . | . | Q | . | . | A | I | G | S | . | . | . | . | . | . | . | . | . | . | . | . | . | . | . | A | . | . | . | . | . | . | . | . | . | . | . | . | . | . | . | . | . | H | . | . | . | . | . | . | . | . | . | A | S | . | . | . | A | K | . | T | . | . | . | D | . | . | . | T | . | . | . | . | S | . | . | . | . | . | . | V | Q | L | . | G | . | S | . |
| Aus0085 | ≥16 | NC_021994.1 | . | . | . | . | . | . | A | . | G | . | . | . | Q | . | . | E | . | T | . | D | . | . | Q | . | . | . | Q | . | . | A | I | G | S | . | . | . | . | . | . | . | . | . | . | . | . | . | . | . | A | . | . | . | . | . | . | . | . | . | . | . | . | . | . | . | . | . | H | . | . | . | . | . | . | . | . | . | A | S | . | . | . | A | K | . | T | . | . | . | D | T | . | . | T | . | . | . | . | S | . | . | . | . | . | . | V | Q | L | . | G | . | S | . |
| E2560 | ≥16 | AHYI01000006.1 | . | . | . | . | . | . | A | . | G | . | . | . | Q | . | . | E | . | T | . | D | . | . | Q | . | . | . | Q | . | . | A | I | G | S | . | . | . | . | . | . | . | . | . | . | . | . | . | . | . | A | . | . | . | . | . | . | . | . | . | . | . | . | . | . | . | . | . | H | . | . | . | . | . | . | . | . | . | A | S | . | . | . | A | K | . | T | . | . | . | D | T | . | . | T | . | . | . | . | S | . | . | . | . | . | . | V | Q | L | . | G | . | S | . |
| U0317 | ≥16 | ABSW01000111.1 | . | . | . | . | . | . | A | . | G | . | . | . | Q | . | . | E | . | T | . | D | . | . | Q | . | . | . | Q | . | . | A | I | G | S | . | . | . | . | . | . | . | . | . | . | . | . | . | . | . | A | . | . | . | . | . | . | . | . | . | . | . | . | . | . | . | . | . | H | . | . | . | . | . | . | . | . | . | A | S | . | . | . | A | K | . | T | . | . | . | D | T | . | . | T | . | . | . | . | S | . | . | . | . | . | . | V | Q | L | . | G | . | S | . |
| D344 | ≥16 | AF362954.1 | . | . | . | . | . | . | A | . | G | . | . | . | Q | . | . | E | . | T | . | D | T | . | Q | . | . | . | Q | . | . | A | I | G | S | . | . | . | . | . | . | . | . | . | . | . | . | . | . | . | A | . | . | . | . | . | . | . | . | . | . | . | . | . | . | . | . | . | . | . | . | . | . | . | . | . | . | . | . | . | . | . | . | T | K | . | T | . | . | . | D | . | . | . | . | . | . | . | . | . | . | . | Y | . | . | E | V | . | . | . | G | A | S | . |
| SN194 | ≥16 | JN208886.1 | . | . | . | . | . | . | A | . | G | . | . | . | Q | . | . | E | . | T | . | D | . | . | Q | . | . | . | Q | . | . | A | I | G | S | . | . | . | R | . | . | . | . | . | . | . | . | . | . | . | A | . | . | . | . | . | . | . | . | . | . | . | . | . | . | D | . | . | . | . | . | . | . | . | . | . | . | . | . | S | D | . | S | | K | L | T | . | . | . | D | . | . | . | . | . | . | . | . | . | . | . | . | . | . | . | V | . | . | . | G | . | S | . |
| SE97F1 | ≥16 | KC479676.1 | . | . | . | . | . | . | A | . | G | . | . | . | Q | . | . | E | . | T | . | D | . | . | Q | . | . | . | Q | . | . | A | I | G | S | . | . | . | . | D | . | . | . | . | . | . | . | . | . | . | A | . | . | . | . | P | . | . | . | . | . | . | . | . | . | D | . | . | . | C | . | . | . | . | . | . | . | K | . | S | . | . | . | A | K | I | T | . | . | . | D | . | . | . | . | . | . | . | . | . | . | . | . | . | . | . | V | . | . | . | G | . | S | . |
| 164306 | ≥16 | JN208882.1 | . | . | . | . | . | . | A | . | G | . | . | . | Q | . | . | E | . | . | . | . | . | . | Q | . | . | . | Q | . | . | A | I | . | S | . | . | . | . | . | . | . | . | . | . | . | . | . | . | . | A | . | . | . | . | . | . | R | . | . | . | . | . | . | S | . | . | . | . | . | . | . | . | . | . | . | . | . | . | . | . | . | . | . | K | . | I | . | . | . | D | . | . | . | . | . | . | . | . | . | . | . | . | . | . | . | . | . | H | . | G | . | . | . |
| H196 | ≥16 | KC479673.1 | . | . | . | . | . | . | A | . | G | . | . | . | Q | . | . | E | . | . | . | . | . | . | Q | . | . | . | Q | . | . | A | I | . | S | . | . | . | . | . | . | . | . | . | . | . | . | . | . | . | A | . | . | . | . | . | . | R | . | . | . | . | . | . | S | . | . | . | . | . | . | . | . | . | . | . | . | . | . | . | . | . | . | . | K | . | I | . | . | . | D | . | . | . | . | . | . | . | . | . | . | . | . | . | . | . | . | . | . | . | G | . | . | . |
| HM1074 | ≥16 | AIUA01000011.1 | . | . | . | . | . | . | A | . | G | . | . | . | Q | . | . | E | . | . | . | . | . | . | Q | . | . | . | Q | . | . | A | I | . | S | . | . | . | . | . | . | . | . | . | . | . | . | . | E | . | A | . | . | . | . | . | . | . | . | . | . | . | . | . | S | . | . | . | . | . | . | . | . | . | . | . | . | . | . | . | . | . | . | . | K | . | I | . | . | . | D | . | . | . | . | . | . | . | . | . | . | . | . | . | . | . | . | . | . | . | G | . | . | . |
| E1636 | ≥16 | ABRY01000062.1 | . | . | . | . | . | . | A | . | G | . | . | . | Q | . | . | E | . | . | . | . | . | . | Q | . | . | . | Q | . | . | A | I | . | S | . | . | . | . | . | . | . | . | . | . | . | . | . | . | . | A | . | . | . | . | . | . | . | . | . | . | . | . | . | S | . | . | . | . | . | . | . | . | . | . | . | . | . | . | . | . | . | . | . | K | . | I | . | . | . | D | . | . | . | . | . | . | . | . | . | . | . | . | . | . | . | . | . | . | . | G | . | . | . |
| HM1072 | ≥16 | AITY01000037.1 | . | . | . | . | . | . | A | . | G | . | . | . | Q | . | . | E | . | . | . | . | . | . | Q | . | . | . | Q | . | . | A | I | . | S | . | . | . | . | . | . | . | . | . | . | . | . | . | . | . | A | . | . | . | . | . | . | . | . | . | . | . | . | . | S | . | . | . | . | . | . | . | . | . | . | . | . | . | . | . | . | . | . | . | K | . | I | . | . | . | D | . | . | . | . | . | . | . | . | . | . | . | . | . | . | . | . | . | . | . | G | . | . | . |
| HM1073 | ≥16 | AITZ01000019.1 | . | . | . | . | . | . | A | . | G | . | . | . | Q | . | . | E | . | . | . | . | . | . | Q | . | . | . | Q | . | . | A | I | . | S | . | . | . | . | . | . | . | . | . | . | . | . | . | . | . | A | . | . | . | . | . | . | . | . | . | . | . | . | . | S | . | . | . | . | . | . | . | . | . | . | . | . | . | . | . | . | . | . | . | K | . | I | . | . | . | D | . | . | . | . | . | . | . | . | . | . | . | . | . | . | . | . | . | . | . | G | . | . | . |
| 9439 | ≥16 | X92687.1 | . | . | . | . | . | . | A | . | G | . | . | . | Q | . | . | E | . | T | . | D | . | . | Q | . | . | . | Q | . | . | A | I | G | S | . | . | . | . | . | . | . | . | . | . | . | . | . | . | . | A | . | . | . | . | . | P | . | . | . | . | . | . | . | . | . | . | . | . | . | I | . | . | . | . | . | . | . | . | . | . | . | . | . | K | . | T | . | . | . | D | . | . | . | . | . | A | . | . | . | L | . | . | . | . | . | V | . | . | . | G | . | S | . |
| EFM-1 | ≥16 | X84861.1 | - | - | - | - | . | . | A | . | G | . | . | . | Q | . | . | E | . | T | . | D | . | . | Q | . | . | . | Q | . | . | A | I | | S | . | . | . | . | . | . | . | . | . | . | . | . | . | . | . | A | . | . | . | . | . | . | . | . | . | . | . | . | . | S | . | . | . | . | . | . | . | . | . | . | . | . | . | . | . | . | . | . | T | K | . | I | . | . | . | D | . | . | . | . | . | . | . | . | . | L | . | . | . | . | . | V | . | . | . | G | . | S | . |
| GEH323_3 | ≥16 | KC479674.1 | . | . | . | . | . | . | A | . | G | . | . | . | Q | . | . | E | . | T | . | D | . | . | Q | . | . | . | Q | . | . | A | I | G | S | . | . | . | . | . | . | . | . | . | . | . | . | . | . | . | A | . | . | . | . | . | . | . | . | . | . | . | . | . | . | D | . | . | H | . | . | . | . | . | . | . | . | . | . | . | . | . | . | A | K | I | T | . | . | . | D | . | . | . | . | . | . | . | . | . | L | . | . | . | . | . | V | . | . | . | G | . | S | . |
| 529940 | ≥16 | JN208883.1 | . | . | . | . | . | . | A | . | G | . | . | . | Q | . | . | E | . | T | . | D | . | . | Q | . | . | . | Q | . | . | A | I | G | S | . | H | . | R | . | S | . | . | . | . | . | . | . | . | . | A | . | . | . | . | . | . | . | . | . | . | . | . | . | . | . | . | . | . | . | . | . | . | . | . | . | . | . | . | . | . | . | . | A | K | L | T | . | . | . | D | . | . | . | . | . | . | . | . | . | L | . | . | . | . | . | V | . | . | . | G | . | S | . |
| GEE4_1 | ≥16 | JN208889.1 | . | . | . | . | . | . | A | . | G | . | . | . | Q | . | . | E | . | T | . | D | . | . | Q | . | . | . | Q | . | . | A | I | G | S | . | . | . | . | . | S | . | . | . | . | . | . | . | . | . | A | . | . | . | E | . | . | . | . | . | . | . | . | . | . | . | . | . | . | . | . | . | . | . | . | . | . | K | . | . | . | . | . | A | K | . | T | . | . | . | D | . | . | . | . | . | . | . | . | . | L | . | . | . | . | . | V | . | . | . | G | . | S | . |
| GEE49_1 | ≥16 | JN208884.1 | . | . | . | . | . | . | A | . | G | . | . | . | Q | . | . | E | . | T | . | D | . | . | Q | . | . | . | Q | . | . | A | I | G | S | . | . | . | . | . | . | . | . | . | . | . | . | . | . | . | A | . | . | . | . | . | . | . | . | . | . | . | . | . | . | D | . | . | . | . | . | . | . | . | . | . | . | K | . | . | . | . | . | A | K | . | T | . | . | . | D | . | . | . | . | . | . | . | . | . | L | . | . | . | . | . | V | . | . | . | G | . | S | . |
| SN449 | ≥16 | JN208887.1 | . | . | . | . | . | . | A | . | G | . | . | . | Q | . | . | E | . | T | . | D | . | . | Q | . | . | . | Q | . | . | A | I | G | S | . | . | . | . | . | . | . | . | . | . | . | . | . | . | . | A | . | . | . | . | . | . | . | . | . | . | . | . | . | . | D | . | . | . | . | . | . | . | . | . | . | . | K | . | . | . | . | . | A | K | . | T | . | . | . | D | . | . | . | . | . | . | . | . | . | L | . | . | . | . | . | V | . | . | . | G | . | S | . |
| Aus0004 | ≥16 | NC_017022.1 | . | . | . | . | . | . | A | . | G | . | . | . | Q | . | . | E | . | T | . | D | . | . | Q | . | . | . | Q | . | . | A | I | G | S | . | . | . | . | . | . | . | . | . | . | . | . | . | . | . | A | . | . | . | . | . | . | . | . | . | . | . | . | . | . | . | . | . | . | . | . | . | . | . | . | . | . | K | . | . | . | . | . | A | K | . | T | . | . | . | D | . | . | . | . | . | . | . | . | . | L | . | . | . | . | . | V | . | . | . | G | . | S | . |
| E1731 | ≥16 | AHXO01000011.1 | . | . | . | . | . | . | A | . | G | . | . | . | Q | . | . | E | . | T | . | D | . | . | Q | . | . | . | Q | . | . | A | I | G | S | . | . | . | . | . | . | . | . | . | . | . | . | . | . | . | A | . | . | . | . | . | . | . | . | . | . | . | . | . | . | . | . | . | . | . | . | . | . | . | . | . | . | K | . | . | . | . | . | A | K | . | T | . | . | . | D | . | . | . | . | . | . | . | . | . | L | . | . | . | . | . | V | . | . | . | G | . | S | . |
| E2883 | ≥16 | AHXX01000007.1 | . | . | . | . | . | . | A | . | G | . | . | . | Q | . | . | E | . | T | . | D | . | . | Q | . | . | . | Q | . | . | A | I | G | S | . | . | . | . | . | . | . | . | . | . | . | . | . | . | . | A | . | . | . | . | . | . | . | . | . | . | . | . | . | . | . | . | . | . | . | . | . | . | . | . | . | . | K | . | . | . | . | . | A | K | . | T | . | . | . | D | . | . | . | . | . | . | . | . | . | L | . | . | . | . | . | V | . | . | . | G | . | S | . |
| E4389 | ≥16 | AHYJ01000006.1, AHYJ01000005.1 | . | . | . | . | . | . | A | . | G | . | . | . | Q | . | . | E | . | T | . | D | . | . | Q | . | . | . | Q | . | . | A | I | G | S | . | . | . | . | . | . | . | . | . | . | . | . | . | . | . | A | . | . | . | . | . | . | . | . | . | . | . | . | . | . | . | . | . | . | . | . | . | . | . | . | . | . | K | . | . | . | . | . | A | K | . | T | . | . | . | D | . | . | . | . | . | . | . | . | . | L | . | . | . | . | . | V | . | . | . | G | . | S | . |
| E1293 | ≥16 | AHWU01000013.1 | . | . | . | . | . | . | A | . | G | . | . | . | Q | . | . | E | . | T | . | D | . | . | Q | . | . | . | Q | . | . | A | I | G | S | . | . | . | . | . | . | . | . | . | . | . | . | . | . | . | A | . | . | . | . | . | . | . | . | . | . | . | . | . | . | . | . | . | . | . | . | . | . | . | . | . | . | K | . | S | . | . | . | A | K | . | T | . | . | . | D | . | . | . | . | . | . | . | . | . | . | . | . | . | . | . | V | . | . | . | G | . | S | . |
| E1644 | ≥16 | AHYG01000012.1 | . | . | . | . | . | . | A | . | G | . | . | . | Q | . | . | E | . | T | . | D | . | . | Q | . | . | . | Q | . | . | A | I | G | S | . | . | . | . | . | . | . | . | . | . | . | . | . | . | . | A | . | . | . | . | . | . | . | . | . | . | . | . | . | . | . | . | . | . | . | . | . | . | . | . | . | . | K | . | S | . | . | . | A | K | . | T | . | . | . | D | . | . | . | . | . | . | . | . | . | . | . | . | . | . | . | V | . | . | . | G | . | S | . |
| GESN71_1 | ≥16 | JN208885.1 | . | . | . | . | . | . | A | . | G | . | . | . | Q | . | . | E | . | T | . | D | . | . | Q | . | . | . | Q | . | . | A | I | G | S | . | . | . | . | . | . | . | . | . | . | . | . | . | . | . | A | V | . | . | . | . | . | . | . | . | . | . | . | . | . | . | . | . | . | . | . | . | . | . | . | . | . | K | . | S | . | . | . | A | K | . | T | . | . | . | D | . | . | . | . | . | . | . | . | . | . | . | . | . | . | . | V | . | . | . | G | . | S | . |
| HPH6 | ≥16 | KC479675.1 | . | . | . | . | . | . | A | . | G | . | . | . | Q | . | . | E | . | T | . | D | . | . | Q | . | . | . | Q | . | . | A | I | G | S | . | . | . | . | . | . | . | . | . | . | . | . | . | . | . | A | . | . | . | . | . | . | . | . | . | . | . | . | . | . | . | . | . | . | . | . | . | . | . | . | . | . | . | . | . | . | . | . | A | K | L | T | . | . | . | D | . | . | . | . | . | . | . | . | . | L | . | . | . | . | . | V | . | . | . | G | . | S | . |
| E0333 | ≥16 | AHWL01000001.1 | . | . | . | . | . | . | A | . | G | . | . | . | Q | . | . | E | . | T | . | D | . | . | Q | . | . | . | Q | . | . | A | I | G | S | . | . | . | . | . | . | . | . | . | . | . | . | . | . | . | A | . | . | . | . | . | . | . | . | . | . | . | . | . | . | . | . | . | . | . | . | . | . | . | . | . | . | . | . | . | . | . | . | T | K | . | T | . | . | . | D | . | . | . | . | . | . | . | . | . | L | . | . | . | . | . | V | . | . | . | G | . | S | . |
| E1162 | ≥16 | ABQJ01000017.1 | . | . | . | . | . | . | A | . | G | . | . | . | Q | . | . | E | . | T | . | D | . | . | Q | . | . | . | Q | . | . | A | I | G | S | . | . | . | . | . | . | . | . | . | . | . | . | . | . | . | A | . | . | . | . | . | . | . | . | . | . | . | . | . | . | . | . | . | . | . | . | . | . | . | . | . | . | . | . | . | . | . | . | T | K | . | T | . | . | . | D | . | . | . | . | . | . | . | . | . | L | . | . | . | . | . | V | . | . | . | G | . | S | . |
| GE70411_2 | ≥16 | JN208888.1 | . | . | . | . | . | . | A | . | G | . | . | . | Q | . | . | E | . | T | . | D | . | . | Q | . | . | . | Q | . | . | A | I | G | S | . | . | . | . | . | . | . | . | . | . | . | . | . | . | . | A | . | . | . | . | . | . | . | . | . | . | . | . | . | . | . | . | . | . | . | . | . | . | . | . | . | . | . | . | . | . | . | . | T | K | . | T | . | . | . | D | . | . | . | . | . | . | . | . | . | L | . | . | . | . | . | V | . | . | . | G | . | S | . |
| HM1071 | ≥16 | AITX01000027.1 | . | . | . | . | . | . | A | . | G | . | . | . | Q | . | . | E | . | T | . | D | . | . | Q | . | . | . | Q | . | . | A | I | G | S | . | . | . | . | . | . | . | . | . | . | . | . | . | . | . | A | . | . | . | . | . | . | . | . | . | . | . | . | . | . | . | . | . | . | . | . | . | . | . | . | . | . | . | . | . | . | . | . | T | K | . | T | . | . | . | D | . | . | . | . | . | . | . | . | . | L | . | . | . | . | . | V | . | . | . | G | . | S | . |
| TX0133a01 | ≥16 | AECJ01000122.1 | . | . | . | . | . | . | A | . | G | . | . | . | Q | . | . | E | . | T | . | D | . | . | Q | . | . | . | Q | . | . | A | I | G | S | . | . | . | . | . | . | . | . | . | . | . | . | . | . | . | A | . | . | . | . | . | . | . | . | . | . | . | . | . | . | . | . | . | . | . | . | . | . | . | . | . | . | . | . | . | . | . | . | T | K | . | T | . | . | . | D | . | . | . | . | . | . | . | . | . | L | . | . | . | . | . | V | . | . | . | G | . | S | . |

| | | | |
|---|---|---|---|
| **TX0133a04** | ≥16 | AEBC01000061.1 | 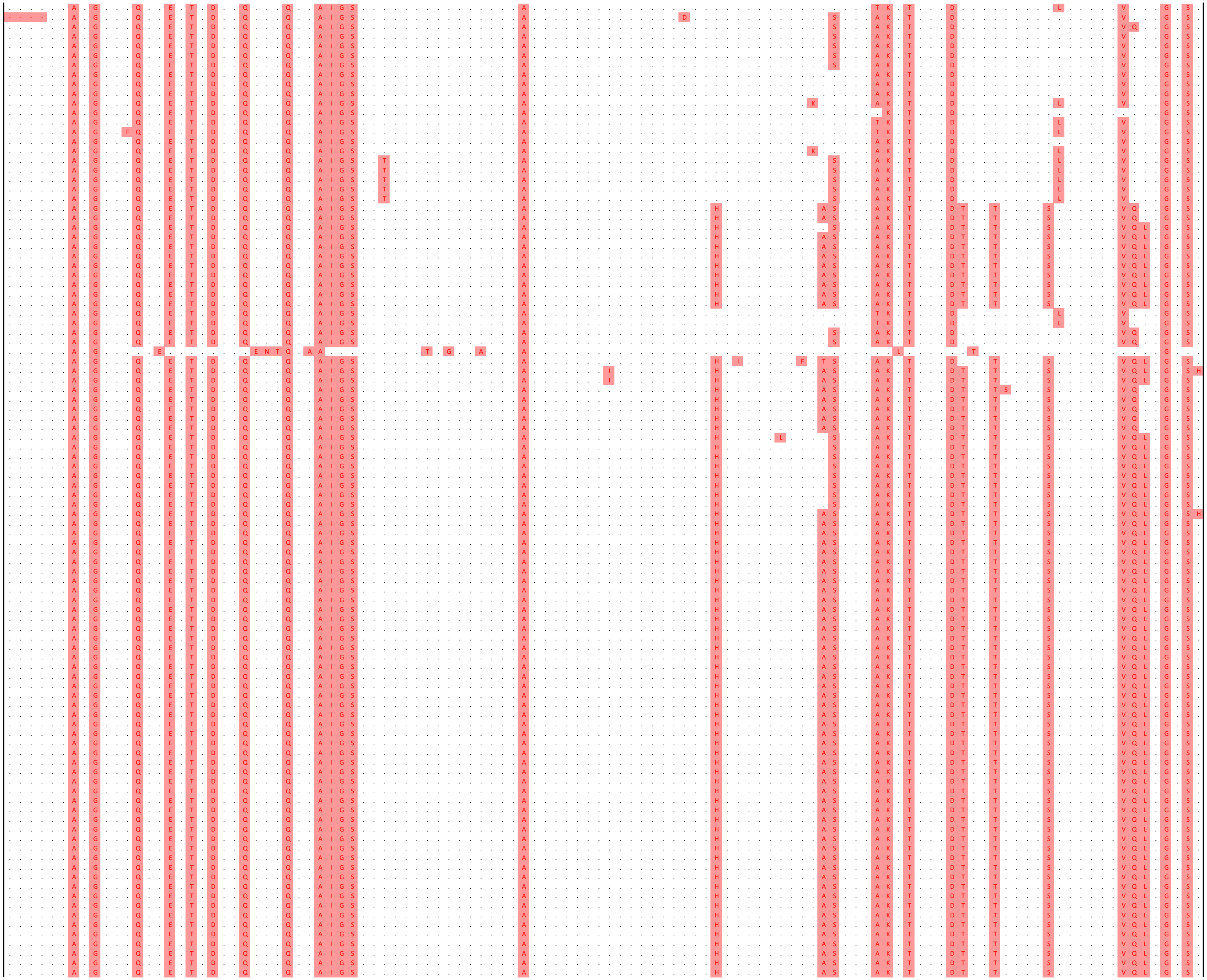 |
| **H80721** | ≥16 | X84862.1 | |
| **1231410** | ≥16 | ACBA01000014.1 | |
| **1231408** | ≥16 | GG688547.1, ACBB01000047.1 | |
| **E1133** | ≥16 | AHWR01000016.1 | |
| **E4452** | ≥16 | AEOU01000130.1 | |
| **E4453** | ≥16 | AEDZ01000364.1 | |
| **E1321** | ≥16 | AHYF01000006.1 | |
| **E1392** | ≥16 | AHWV01000002.1 | |
| **E1679** | ≥16 | ABSC01000284.1 | |
| **NEF1** | ≥16 | AXOK01000016.1, AXOK01000017.1 | |
| **DO** | 16 | NC_017960.1 | |
| **TX0133B** | 16 | AECI01000046.1 | |
| **TX0133A** | 32 | AECH01000021.1 | |
| **E1904** | 32 | AHXQ01000004.1 | |
| **ERV99** | >32/R | AMAQ01 | |
| **ERV26** | >32/R | AMAT01 | |
| **ERV28** | >32/R | MTRY00000000 | |
| **ERV30** | >32/R | MTRW00000000 | |
| **ERV34** | >32/R | MJFW00000000 | |
| **ERV35** | >32/R | MJFX00000000 | |
| **ERV218** | 64 | MJFJ00000000 | |
| **ERV223** | 64 | MJFL00000000 | |
| **2043** | 64 | Bhavarth | |
| **2036** | 64 | Bhavarth | |
| **2053** | 64 | Bhavarth | |
| **ERV174** | 64 | MJFC00000000 | |
| **ERV175** | 64 | MJFD00000000 | |
| **ERV177** | 64 | MJFE00000000 | |
| **ERV180** | 64 | MJFF00000000 | |
| **ERV212** | 64 | MTRX00000000 | |
| **ERV214** | 64 | MJFI00000000 | |
| **TX0082** | 64 | AEBU01000140.1 | |
| **TX0133C** | 64 | AEBG01000194.1 | |
| **ERV38** | 64 | AMAS01 | |
| **ERV69** | 64 | AMAR01 | |
| **505** | >64/R | AMBL01000029.1, AMBL01000072.1 | |
| **MB871** | >64/R | Bhavarth | |
| **UVA5** | >64/R | Bhavarth | |
| **MB021** | >64/R | Bhavarth | |
| **ERV98** | >64/R | MJFY00000000 | |
| **C621** | >64/R | AMBB01 | |
| **ERV102** | >64/R | AMAX01 | |
| **ERV89** | >64/R | MTRT00000000 | |
| **MB069** | >64/R | Bhavarth | |
| **C11527** | >64/R | Bhavarth | |
| **1979** | >64/R | Bhavarth | |
| **1987** | >64/R | Bhavarth | |
| **2019** | >64/R | Bhavarth | |
| **2020** | >64/R | Bhavarth | |
| **ERV275** | >64/R | MJFS00000000 | |
| **ERV280** | >64/R | MJFU00000000 | |
| **1996** | >64/R | Bhavarth | |
| **ERV251** | >64/R | MJFP00000000 | |
| **UVA8** | >64/R | Bhavarth | |
| **UVA7** | >64/R | Bhavarth | |
| **S11491** | >64/R | Bhavarth | |
| **MB662** | >64/R | Bhavarth | |
| **MB613** | >64/R | Bhavarth | |
| **MB460** | >64/R | Bhavarth | |
| **MB442** | >64/R | Bhavarth | |
| **MB278** | >64/R | Bhavarth | |
| **MB119** | >64/R | Bhavarth | |
| **MB1054** | >64/R | Bhavarth | |
| **2045** | >64/R | Bhavarth | |
| **2033** | >64/R | Bhavarth | |
| **2030** | >64/R | Bhavarth | |
| **2027** | >64/R | Bhavarth | |
| **2008** | >64/R | Bhavarth | |
| **1999** | >64/R | Bhavarth | |
| **1994** | >64/R | Bhavarth | |
| **1992** | >64/R | Bhavarth | |
| **1991** | >64/R | Bhavarth | |
| **1982** | >64/R | Bhavarth | |
| **1977** | >64/R | Bhavarth | |
| **1963** | >64/R | Bhavarth | |
| **1958** | >64/R | Bhavarth | |
| **1953** | >64/R | Bhavarth | |
| **1954** | >64/R | Bhavarth | |
| **1955** | >64/R | Bhavarth | |
| **1975** | >64/R | Bhavarth | |
| **1985** | >64/R | Bhavarth | |
| **1988** | >64/R | Bhavarth | |
| **1998** | >64/R | Bhavarth | |
| **2003** | >64/R | Bhavarth | |
| **2007** | >64/R | Bhavarth | |
| **2009** | >64/R | Bhavarth | |
| **2022** | >64/R | Bhavarth | |
| **2040** | >64/R | Bhavarth | |
| **2050** | >64/R | Bhavarth | |
| **2056** | >64/R | Bhavarth | |
| **511** | >64/R | AMBH01000038.1 | |
| **513** | >64/R | AMBG01000061.1 | |
| **514** | >64/R | AMBF01000038.1 | |
| **515** | >64/R | AMBE01000026.1 | |
| **E422** | >64/R | AMAZ01000004.1 | |
| **ERV115** | >64/R | MTRV00000000 | |
| **ERV118** | >64/R | MTRU00000000 | |
| **ERV133** | >64/R | MJEY00000000 | |
| **ERV155** | >64/R | MJEZ00000000 | |
| **ERV157** | >64/R | MJFA00000000 | |
| **ERV161** | >64/R | AMAW01 | |

| | | | |
|---|---|---|---|
| **ERV165** | >64/R | AMAV01 | ......A.G...Q..E.T.D..Q...Q..AIGS...............A.................H.........AS...AK.T...DT..T....S......VQL.G.S. |
| **ERV168** | >64/R | AMAU01 | ......A.G...Q..E.T.D..Q...Q..AIGS...............A.................H.........AS...AK.T...DT..T....S......VQL.G.S. |
| **ERV172** | >64/R | MJFB00000000 | ......A.G...Q..E.T.D..Q...Q..AIGS...............A.................H.........AS...AK.T...DT..T....S......VQL.G.S. |
| **ERV190** | >64/R | MJFG00000000 | ......A.G...Q..E.T.D..Q...Q..AIGS...............A.................H.........AS...AK.T...DT..T....S......VQL.G.S. |
| **ERV196** | >64/R | MJFH00000000 | ......A.G...Q..E.T.D..Q...Q..AIGS...............A.................H.........AS...AK.T...DT..T....S......VQL.G.S. |
| **ERV220** | >64/R | MJFK00000000 | ......A.G...Q..E.T.D..Q...Q..AIGS...............A.................H.........AS...AK.T...DT..T....S......VQL.G.S. |
| **ERV231** | >64/R | MJFM00000000 | ......A.G...Q..E.T.D..Q...Q..AIGS...............A.................H.........AS...AK.T...DT..T....S......VQL.G.S. |
| **ERV237** | >64/R | MJFN00000000 | ......A.G...Q..E.T.D..Q...Q..AIGS...............A.................H.........AS...AK.T...DT..T....S......VQL.G.S. |
| **ERV242** | >64/R | MJFO00000000 | ......A.G...Q..E.T.D..Q...Q..AIGS...............A.................H.........AS...AK.T...DT..T....S......VQL.G.S. |
| **ERV263** | >64/R | MJFQ00000000 | ......A.G...Q..E.T.D..Q...Q..AIGS...............A.................H.........AS...AK.T...DT..T....S......VQL.G.S. |
| **ERV270** | >64/R | MJFR00000000 | ......A.G...Q..E.T.D..Q...Q..AIGS...............A.................H.........AS...AK.T...DT..T....S......VQL.G.S. |
| **ERV281** | >64/R | MJFV00000000 | ......A.G...Q..E.T.D..Q...Q..AIGS...............A.................H.........AS...AK.T...DT..T....S......VQL.G.S. |
| **M11518** | >64/R | Bhavarth | ......A.G...Q..E.T.D..Q...Q..AIGS...............A.................H.........AS...AK.T...DT..T....S......VQL.G.S. |
| **MB020** | >64/R | Bhavarth | ......A.G...Q..E.T.D..Q...Q..AIGS...............A.................H.........AS...AK.T...DT..T....S......VQL.G.S. |
| **MB167** | >64/R | Bhavarth | ......A.G...Q..E.T.D..Q...Q..AIGS...............A.................H.........AS...AK.T...DT..T....S......VQL.G.S. |
| **MB263** | >64/R | Bhavarth | ......A.G...Q..E.T.D..Q...Q..AIGS...............A.................H.........AS...AK.T...DT..T....S......VQL.G.S. |
| **MB271** | >64/R | Bhavarth | ......A.G...Q..E.T.D..Q...Q..AIGS...............A.................H.........AS...AK.T...DT..T....S......VQL.G.S. |
| **MB304** | >64/R | Bhavarth | ......A.G...Q..E.T.D..Q...Q..AIGS...............A.................H.........AS...AK.T...DT..T....S......VQL.G.S. |
| **MB407** | >64/R | Bhavarth | ......A.G...Q..E.T.D..Q...Q..AIGS...............A.................H.........AS...AK.T...DT..T....S......VQL.G.S. |
| **MB443** | >64/R | Bhavarth | ......A.G...Q..E.T.D..Q...Q..AIGS...............A.................H.........AS...AK.T...DT..T....S......VQL.G.S. |
| **MB663** | >64/R | Bhavarth | ......A.G...Q..E.T.D..Q...Q..AIGS...............A.................H.........AS...AK.T...DT..T....S......VQL.G.S. |
| **P1123** | >64/R | JH810542.1, AMAP01000100.1 | ......A.G...Q..E.T.D..Q...Q..AIGS...............A.................H.........AS...AK.T...DT..T....S......VQL.G.S. |
| **P1139** | >64/R | AMAN01 | ......A.G...Q..E.T.D..Q...Q..AIGS...............A.................H.........AS...AK.T...DT..T....S......VQL.G.S. |
| **P1140** | >64/R | JH808980.1, AMAM01000021.1 | ......A.G...Q..E.T.D..Q...Q..AIGS...............A.................H.........AS...AK.T...DT..T....S......VQL.G.S. |
| **P1986** | >64/R | JH808239.1, AMAK01000017.1 | ......A.G...Q..E.T.D..Q...Q..AIGS...............A.................H.........AS...AK.T...DT..T....S......VQL.G.S. |
| **R446** | >64/R | JH808025.1, AMAJ01000023.1 | ......A.G...Q..E.T.D..Q...Q..AIGS...............A.................H.........AS...AK.T...DT..T....S......VQL.G.S. |
| **R494** | >64/R | AMAI01000028.1 | ......A.G...Q..E.T.D..Q...Q..AIGS...............A.................H.........AS...AK.T...DT..T....S......VQL.G.S. |
| **R496** | >64/R | AMAH01000068.1 | ......A.G...Q..E.T.D..Q...Q..AIGS...............A.................H.........AS...AK.T...DT..T....S......VQL.G.S. |
| **R497** | >64/R | AMAG01000075.1 | ......A.G...Q..E.T.D..Q...Q..AIGS...............A.................H.........AS...AK.T...DT..T....S......VQL.G.S. |
| **R499** | >64/R | AMAF01000040.1 | ......A.G...Q..E.T.D..Q...Q..AIGS...............A.................H.........AS...AK.T...DT..T....S......VQL.G.S. |
| **S447** | >64/R | JH806868.1, AMAD01000018.1 | ......A.G...Q..E.T.D..Q...Q..AIGS...............A.................H.........AS...AK.T...DT..T....S......VQL.G.S. |
| **T21567** | >64/R | Bhavarth | ......A.G...Q..E.T.D..Q...Q..AIGS...............A.................H.........AS...AK.T...DT..T....S......VQL.G.S. |
| **UVA10** | >64/R | Bhavarth | ......A.G...Q..E.T.D..Q...Q..AIGS...............A.................H.........AS...AK.T...DT..T....S......VQL.G.S. |
| **V689** | >64/R | AMAC01000037.1 | ......A.G...Q..E.T.D..Q...Q..AIGS...............A.................H.........AS...AK.T...DT..T....S......VQL.G.S. |
| **504** | >64/R | AMBM01000063.1 | ......A.G...Q..E......Q...Q..AI.S...............A..K..........S......S............K.I...D...................G... |
| **506** | >64/R | JH813945.1, AMBK01000250.1 | ......A.G...Q..E.T.D..Q...Q..AIGS...............A..........................K.....AK.T...D.........L.....V...G.S. |
| **510** | >64/R | AMBI01000132.1 | ......A.G...Q..E.T.D..Q...Q..AIGS...............A..........................K.....AK.T...D.........L.....V...G.S. |
| **C1904** | >64/R | AMBD01 | ......A.G...Q..E.T.D..Q...Q..AIGS...............A..........................K.....AK.T...D.........L.....V...G.S. |
| **C497** | >64/R | AMBC01000119.1 | ......A.G...Q..E.T.D..Q...Q..AIGS...............A..........................K.....AK.T...D.........L.....V...G.S. |
| **ERV279** | >64/R | MJFT00000000 | ......A.G...Q..E.T.D..Q...Q..AIGS...............A..........................K.....AK.T...D.........L.....V...G.S. |
| **MB197** | >64/R | Bhavarth | ......A.G...Q..E.T.D..Q...Q..AIGS...............A..........................K.....AK.T...D.........L.....V...G.S. |
| **P1137** | >64/R | AMAO01000169.1 | ......A.G...Q..E.T.D..Q...Q..AIGS...............A..........................K.....AK.T...D.........L.....V...G.S. |
| **P1190** | >64/R | JH808818.1, AMAL01000087.1 | ......A.G...Q..E.T.D..Q...Q..AIGS...............A..........................K.....AK.T...D.........L.....V...G.S. |
| **E417** | >64/R | JH813696.1, AMBA01000202.1 | ......A.G...Q..E.T.D..Q...Q..AIGS...............A............................SD..AK.T...D.........L.....V...G.S. |
| **ERV127** | >64/R | MJEX00000000 | ......A.G...Q..E.T.D..Q...Q..AIGS...............A............................SD..AK.T...D.........L.....V...G.S. |
| **503** | >64/R | JH809465.1, AMBN01000040.1 | ......A.G...Q..E.T.D..Q...Q..AIGS...............A............................S...AK.T...D.........L.....V...G.S. |
| **R501** | >64/R | AMAE01000246.1 | ......A.G...Q..E.T.D..Q...Q..AIGS...............A............................S...AK.T...D...............VQ..G.S. |
| **509** | >64/R | AMBJ01000257.1 | ......A.G...Q..E.T.D..Q...Q..AIGS...............A............................S...AK.T...D...............V...G.S. |
| **ERV113** | >64/R | MJEW00000000 | ......A.G...Q..E.T.D..Q...Q..AIGS...............A............................S...AK.T...D...............V...G.S. |
| **1231502** | 128 | ACAX01 | ......A.G...Q..E.T.D..Q...Q..AIGS...............A.................H.........AS...AK.T...DT..T....S......VQL.G.S. |
| **1230933** | 128 | ACAS01 | ......A.G...Q..E.T.D..Q...Q..AIGS...............A..........................K.....AK.T...D.........L.....V...G.S. |
| **E2134** | 128 | AHXU01 | ......A.G...Q..E.T.D..Q...Q..AIGS...............A..........................K.S...AK.T...D...............V...G.S. |
| **E1626** | 128 | AHXI01 | ......A.G...Q..E.T.D..Q...Q..AIGS...............A............................SD..AK.T...D.........L.....V...G.S. |
| **C68** | 128 | LPUE01 | ......A.G...Q..E.T.D..Q...Q..AIGS...............A............................S...AKLT...D...............V...G.S. |
| **ERV1** | 256 | AMAY01 | ......A.G...Q..E.T.D..Q...Q..AIGS...............A............................SD..AK.T...D.........L.....V...G.S. |

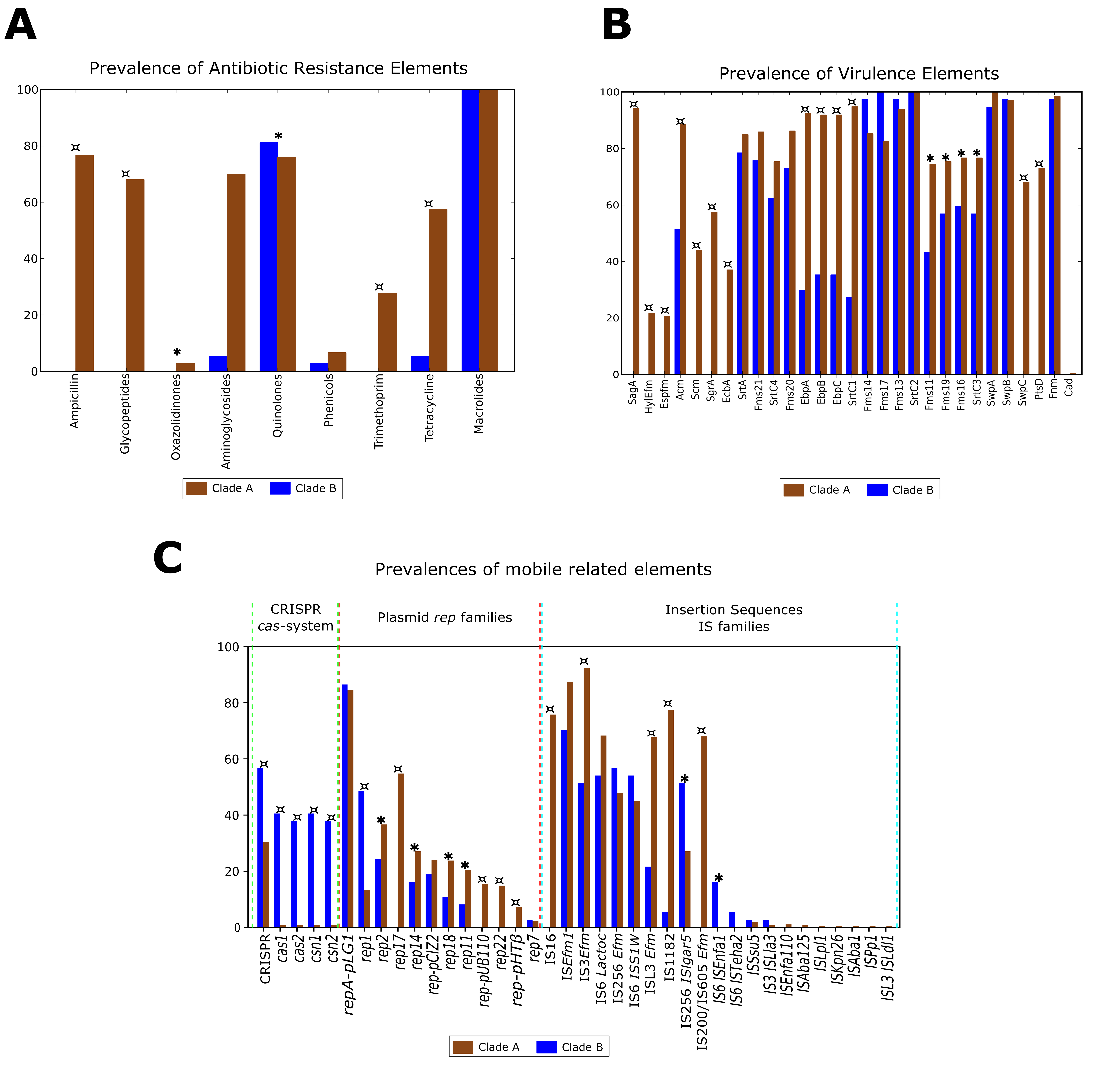

**Supplementary Figure 1.** (A) Prevalences of genetic elements associated to resistance against the groups of antibiotics on each of the clades. Ampicillin resistance was predicted based on a random forest model over the sequence of the PBP5 protein of susceptible and resistance isolates. (B) Prevalences of mobile genetic elements on each of the clades,separated by groups in CRISPR *cas*-system, plasmid *rep* families and insertion sequences. (C) Prevalences of virulence associated elements on each of the clades.

* p-value ≤ 0.05
p-value ≤ 0.001

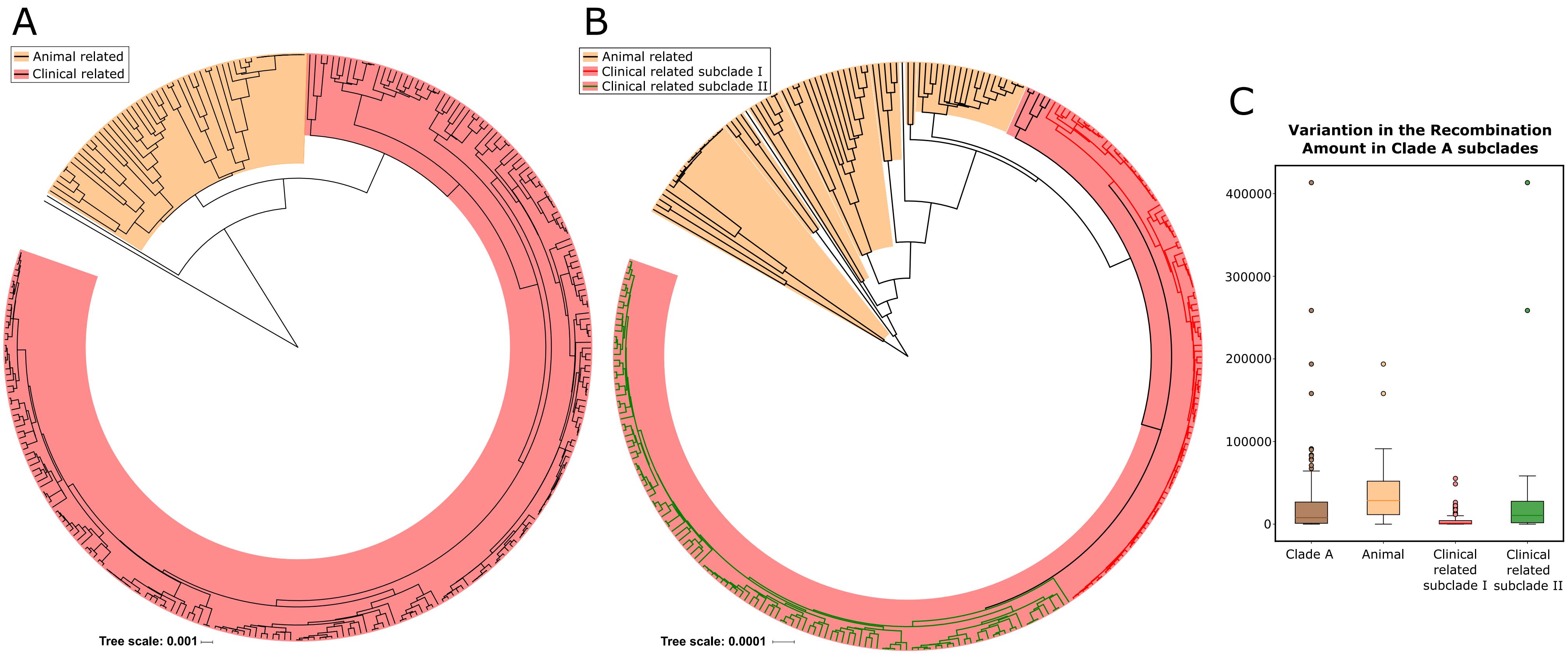

**Supplemental Figure 2.** Comparison of maximum clade credibility trees without excluding recombinant regions (A) and excluding recombinant regions (B). Both trees show the subclades/branches related to animal sources (orange background) and clinical sources (red background). (C) Box and whiskers plot of the amount of recombination in the genomes belonging to the different branches/subclades in the clade A, based on the recombination regions defined by ClonalFrameML, colored dots represent outliers within each group.

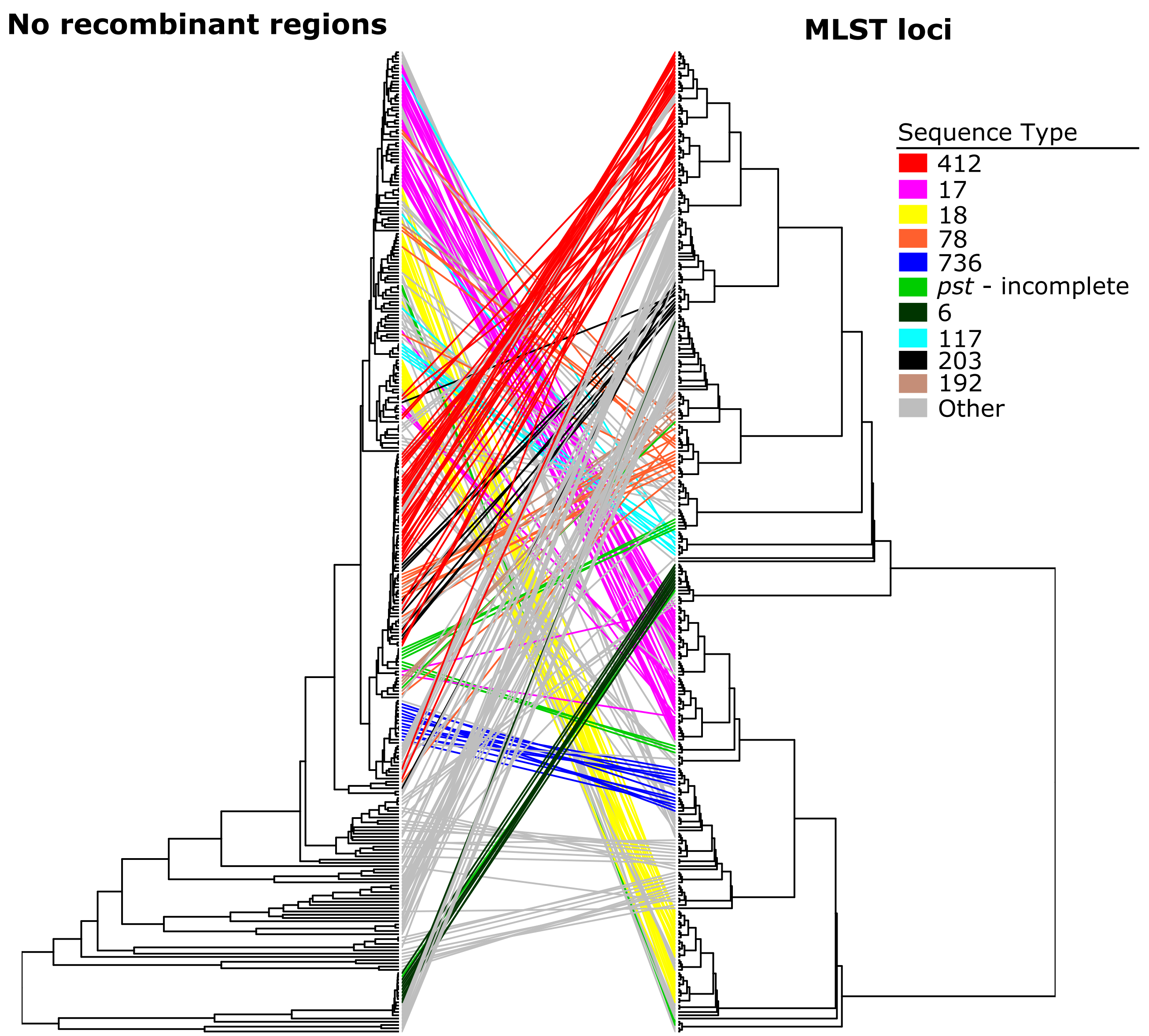

**Supplementary figure 3**. Tanglegram of the phylogenetic reconstructions of Clade A genomes based on the non-recombinant regions (left) against the MLST loci (right), colored lines in the middle connect the branches of the same genome between the two trees.

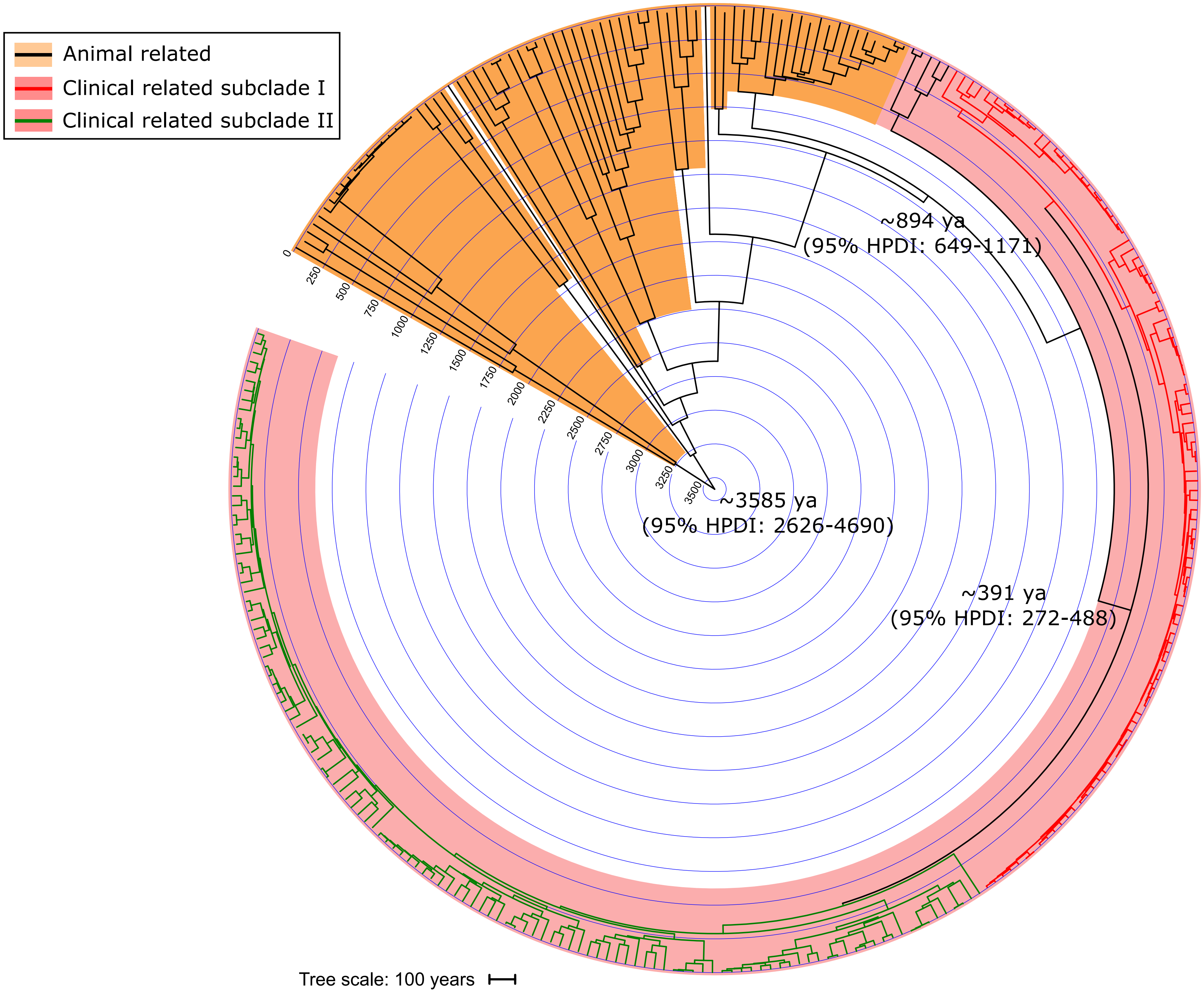

**Supplementary Figure 3**. Maximum Credibility Tree of the molecular clock analysis of clade A genomes. Each blue ring represents a distance of 250 years across branch lengths. At splits between major subclades is the estimated divergence time between them and the 95% High Probability Density Interval (HPDI).
